# A Tripartite Map of the Ventromedial Prefrontal Cortex

**DOI:** 10.64898/2026.06.18.732791

**Authors:** Guoqiu Chen, Donghui Song, Mingxue Fu, Siyi Li, Mingzhe Zhang, Zaixu Cui, Mingrui Xia, Lianglong Sun, Yong He, Ting Xu, Xi Yu, Yinyin Zang, Jingfeng Zhou, Kai Zhang, Shaozheng Qin, Haroon Popal, Zeynep M Saygin, David E Osher, Ingrid R. Olson, Matthew F.S. Rushworth, Yin Wang

## Abstract

The ventromedial prefrontal cortex (VMPFC) has been repeatedly implicated in affect, valuation, and social cognition, yet how these diverse functions are organized within a single cortical territory has remained unresolved. Here, we integrate large-scale meta-analysis, individual-level task fMRI, artificial neural-network encoding models, and multimodal connectivity analyses to reveal the internal functional architecture of the human VMPFC. Across four complementary studies, we identify a robust tripartite organization along the anterior–posterior axis, comprising posterior affective, middle valuation, and anterior social functional motifs. Connectivity fingerprinting demonstrates that each motif is preferentially embedded within distinct large-scale brain networks, providing a mechanistic account of VMPFC functional specialization. This organization is reproducible at the level of individual subjects, generalizes to naturalistic stimuli, extends across development, and shows cross-species correspondence with non-human primates and multiple neurobiological markers. Together, these findings resolve a long-standing organizational question and establish a biologically grounded framework for interpreting VMPFC function.

## Introduction

The human ventromedial prefrontal cortex (VMPFC) has long been a central focus of neuroscience owing to its involvement in emotion, decision-making, and social behavior^1^. Early lesion studies, including the historical case of Phineas Gage, revealed its importance for personality and affective regulation^2^, while subsequent neuroimaging work identified the VMPFC as one of the most metabolically active and evolutionarily expanded regions of the human cortex^3,4^. Positioned along the cortical midline and densely interconnected with distributed systems—including the default mode, limbic, frontoparietal, and salience networks—the VMPFC occupies a central hub in large-scale brain organization^5^. Functionally, it has been implicated across a wide range of psychological phenomena, including affective regulation, valuation, self-referential processing, social inference, memory, and autonomic control^1^, and its dysfunction is associated with multiple psychiatric conditions such as depression, addiction, psychopathy, and autism^6^. Despite decades of intensive research, however, a principled account explaining why this single region consistently recurs across such diverse literatures remains lacking.

Progress toward identifying the functional architecture of the VMPFC has been hindered by several factors. First, the term “VMPFC” does not correspond to a single anatomically circumscribed area but spans multiple cytoarchitectonic subdivisions whose boundaries remain debated^7–9^. Second, the region exhibits pronounced functional heterogeneity, participating in affective, motivational, and social processes that are often investigated in isolation^1^. Third, substantial interindividual variability in sulcal morphology shifts the spatial localization of functional peaks, complicating group-level inference^10,11^. Finally, the ventral prefrontal cortex is particularly vulnerable to susceptibility artifacts in conventional fMRI, leading to signal dropout and spatial distortion unless acquisition is carefully optimized^12–14^. Together, these factors have contributed to a fragmented literature in which distinct functional claims coexist without being integrated into a coherent organizational framework.

In other complex association cortices, such fragmentation has been addressed through data-driven parcellation frameworks that explicitly link anatomy, connectivity, and function into reproducible subregional maps. By identifying spatially contiguous subregions with distinct connectivity and functional profiles, such frameworks help resolve ambiguous boundaries, accommodate functional heterogeneity, and provide a common reference space that is robust to individual variability and methodological differences. Applied to regions such as the amygdala, insula, anterior cingulate cortex, and thalamus, parcellation approaches have clarified longstanding theoretical debates, standardized cross-study comparisons, and accelerated translational research^15–20^. Despite these advances, a comparable multimodal, data-driven functional atlas for the VMPFC has not yet been established.

Although theories of VMPFC function vary widely, prior work has repeatedly suggested that its functional heterogeneity may reflect multiple interacting processes related to affect, valuation, and social cognition^1,6,21^. Meta-analytic studies have reported partially dissociable activation patterns associated with these processes^6,22^, and connectivity studies have described preferential coupling of anterior VMPFC (BA 10) with social brain networks^23^, middle VMPFC (BA 32) with reward-related circuitry^24,25^, and posterior VMPFC (BA 25) with limbic and autonomic systems^22,26^. However, these distinctions have largely emerged from isolated modalities, task paradigms, or analytic traditions, and thus remain conceptually suggestive rather than organizationally definitive. In particular, it has not been established whether such recurring dissociations reflect a stable and reproducible organizational principle of the VMPFC, or instead arise from context-dependent task demands, averaging across heterogeneous individuals, or methodological idiosyncrasies. Critically, prior work has not jointly tested whether putative VMPFC subdivisions are reproducible at the level of individual brains, generalize beyond experimenter-defined tasks to naturalistic stimulus spaces, and are grounded in identifiable large-scale network mechanisms.

Here, we address this gap by testing whether a convergent, tripartite functional organization constitutes a stable and biologically meaningful description of the human VMPFC. Throughout this paper, we use the term *functional motif* to denote a recurrent and dominant pattern of functional organization that preferentially weights distinct psychological processes while remaining embedded within a continuous and integrative circuit architecture, rather than a rigid, domain-specific module. Leveraging a set of complementary, independently grounded approaches—including meta-analytic co-activation modeling, individual-level task-based fMRI, artificial neural-network (ANN) encoding models, connectivity fingerprinting, and lifespan and non-human primate datasets—we address four central questions: (1) What are the dominant functional motifs of the VMPFC, and how are they spatially organized at a coarse scale? (2) Can this organization be reproducibly localized at the level of individual brains using both empirical and computational models of neural response? (3) What large-scale network principles give rise to motif-specific functional specialization within the VMPFC? (4) Does this organizational scheme generalize across development and species? By integrating convergent evidence across scales, modalities, and populations, we delineate anterior social, middle valuation, and posterior affective motifs of the VMPFC, demonstrate how their functional specializations arise from distinct network embeddings, and establish their stability across development and evolution. This tripartite motif-based framework provides a principled reference map for interpreting the VMPFC’s roles in cognition, emotion, and psychopathology, and offers a foundation for future mechanistic and translational research^1,6,21^.

## Results

### Study 1 | Characterizing the functional diversity of the VMPFC

To delineate the dominant functional associations of the ventromedial prefrontal cortex (VMPFC), we first performed large-scale reverse-inference and co-activation meta-analyses using the Neurosynth database (n = 14,371 fMRI studies). Reverse-inference mapping identified the top 15 functional terms most frequently associated with VMPFC activation (Fig. 1a). These terms exhibited an apparent semantic structure and were provisionally organized into three broad categories—social, valuation, and affect—suggesting intrinsic functional heterogeneity (see the full list of top 50 functional terms in Supplementary Table 1).

**Figure 1.**
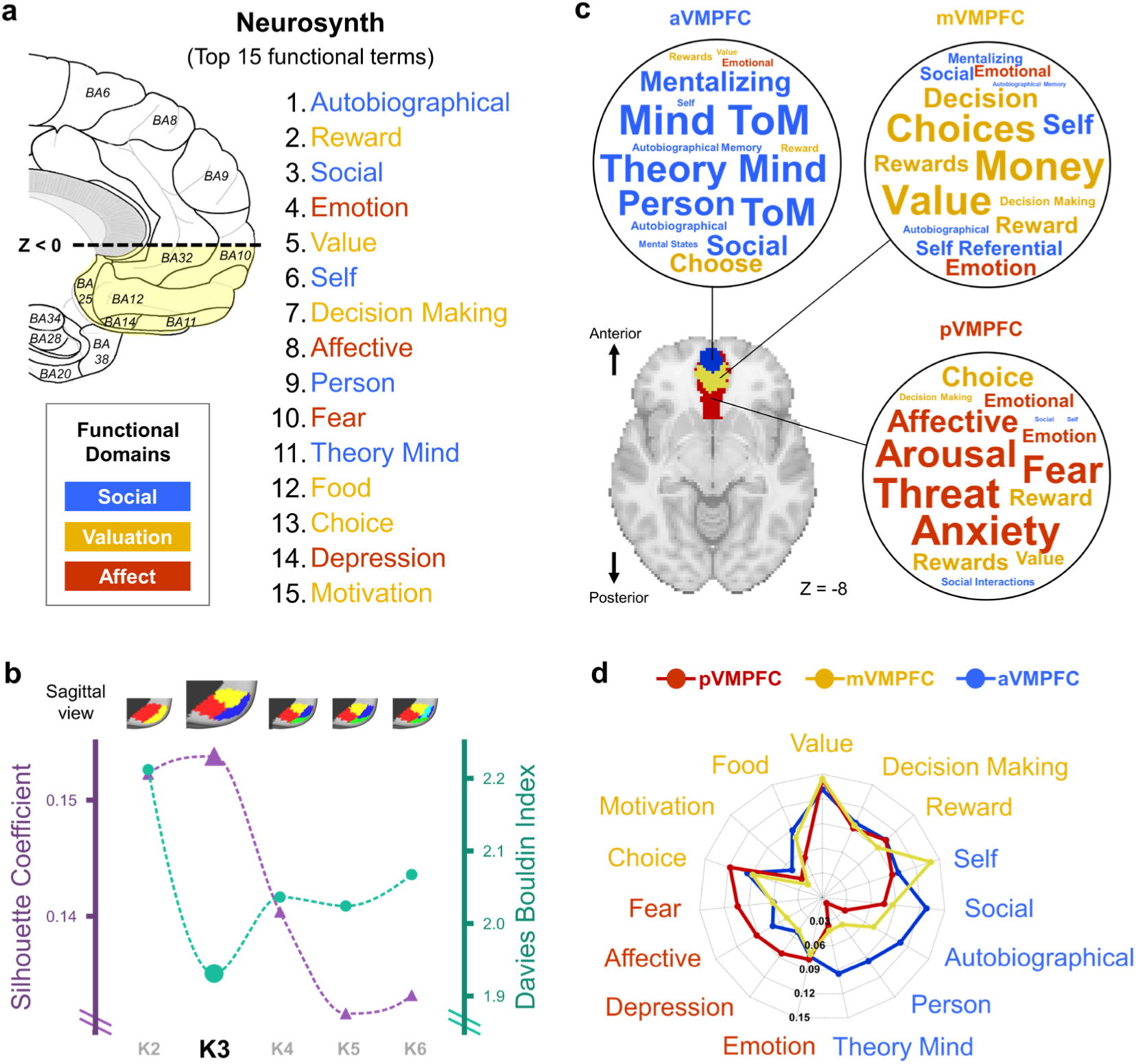
Meta-analytic parcellation and functional decoding reveal a tripartite organization of the human VMPFC into three functional motifs in Study 1. **a,** reverse-inference analysis ranks the top 15 functional terms associated with the VMPFC in the Neurosynth database; terms are color-coded by domain (blue=social; yellow=valuation; red=affect). The left drawing shows the VMPFC definition used in this study (z<0, adopted from an authoritative review article^1^) and its Brodmann coverage. **b**, MACM-based clustering of VMPFC co-activation patterns across the whole brain for K=2–6 (top, sagittal views of the VMPFC parcels). Model-selection metrics (i.e., silhouette coefficient, higher is better; Davies–Bouldin index, lower is better) jointly identify K=3 as the optimal solution. **c,** For each MACM parcel, word clouds display the most diagnostic reverse-inference terms, illustrating three motif-dominant profiles along the posterior-anterior axis: an affective/emotion motif (posterior VMPFC), a valuation/decision motif (middle VMPFC), and a social/mentalizing motif (anterior VMPFC). See its sagittal view in panel b. **d,** Radar plots quantify functional decoding (reverse-inference probabilities) for each parcel across the top 15 functional terms, demonstrating motif-dominant organization along the posterior-to-anterior axis (posterior VMPFC = affective/emotion; middle VMPFC = valuation/decision; anterior VMPFC = social/mentalizing). Similar results can be found in Extended Data Fig. 6d. Colors are consistent across panels.

To test whether this semantic organization reflects a robust data-driven structure, we performed additional term-clustering analyses in two independent representational spaces (Extended Data Fig. 9). First, we embedded the Neurosynth terms using a pretrained language model and applied hierarchical clustering in semantic space. Second, we clustered the same terms using their whole-brain Neurosynth association maps, excluding voxels within the VMPFC mask to avoid circularity with the VMPFC parcellation. Across both the original top-15 terms and an expanded set of 50 terms, the two approaches recovered a highly similar three-cluster structure. Finally, an independent language model was used to assign functional labels to the recovered clusters, yielding labels semantically equivalent to affect, valuation, and social cognition with high consistency across repeated runs (Extended Data Fig. 9b). These results provide quantitative support for the tripartite functional interpretation of the VMPFC motifs while reducing reliance on investigator-defined categorization.

We next applied meta-analytic co-activation modeling (MACM)^27^ to test whether this functional heterogeneity corresponded to separable subregions within the VMPFC. By clustering whole-brain co-activation patterns for the VMPFC across studies, we generated parcellation solutions from K = 2 to K = 6 (Fig. 1b). Quantitative model-selection criteria converged on a three-parcel solution (K = 3), which simultaneously maximized between-cluster separability (Silhouette coefficient) and minimized within-cluster redundancy (Davies–Bouldin index), indicating an optimal balance between parsimony and functional specificity.

To evaluate the robustness of this tripartite solution, we conducted additional analyses varying both voxel-selection criteria and clustering algorithms (Supplementary Fig. 3a). Repeating K-means clustering across different minimum-study inclusion thresholds (≥100, ≥150, and ≥200 studies per voxel) consistently recovered highly similar tripartite solutions, with median Dice similarities exceeding 0.90. Moreover, Spectral Clustering and Agglomerative Clustering converged on comparable posterior–middle–anterior organizations despite relying on fundamentally different optimization principles. These findings indicate that the observed tripartite structure is a stable property of the underlying co-activation architecture rather than an artifact of initialization, voxel-selection criteria, or clustering algorithm. Thus, the tripartite solution reflects a consensus structure recovered across multiple analytical frameworks rather than a specific outcome of K-means optimization.

To further evaluate whether the tripartite organization could be driven by biases in the composition of the underlying literature, we quantified the distribution of studies across functional domains and replicated the analysis in an independent meta-analytic database (Supplementary Fig. 4). Within Neurosynth, affective-, valuation-, and social-related studies were represented at broadly comparable frequencies both across the full database and within the subset of studies contributing to the VMPFC meta-analysis (Supplementary Fig. 4a). Because Neurosynth does not provide manually curated task annotations, we repeated the analysis using the independent BrainMap^28,29^ database (N = 4,341 studies), which contains expert-coded experimental paradigms. Re-running the complete MACM pipeline on BrainMap recovered a highly similar tripartite organization (Dice = 0.77 relative to Neurosynth), and cluster-validity metrics again supported a three-cluster solution (Supplementary Fig. 4b). Domain-level counts remained balanced in BrainMap (Supplementary Fig. 4c), and no individual experimental paradigm accounted for more than 13.5% of VMPFC-activating studies (Supplementary Fig. 4d). Together, these findings indicate that the tripartite VMPFC organization is unlikely to be explained by overrepresentation of particular functional domains or task types within the underlying literature.

To functionally characterize each parcel, we performed reverse-inference decoding of the meta-analytic activation map associated with each cluster. Word-cloud visualizations and quantitative radar plots (Fig. 1c–d) revealed a consistent posterior–anterior ordering of dominant functional associations across the three parcels. Specifically, the posterior VMPFC (pVMPFC) was preferentially associated with affect-related terms such as fear, arousal, and threat; the middle VMPFC (mVMPFC) with valuation- and decision-related terms such as value, money, and choice; and the anterior VMPFC (aVMPFC) with social and self-referential terms such as theory of mind, mentalizing, and autobiographical processing. This ordered pattern indicates systematic differentiation in functional weighting across spatially discrete VMPFC subregions.

Importantly, comparison across alternative parcellation resolutions (K = 2–6; Extended Data Fig. 2) showed that this tripartite functional structure is preserved across resolutions: posterior parcels consistently exhibited affect-related dominance, middle parcels valuation-related dominance, and anterior parcels social-related dominance. Lower-dimensional solutions (K = 2) merged adjacent motifs into broader, mixed profiles, whereas higher-dimensional solutions (K ≥ 4) subdivided these motifs into smaller parcels that retained their dominant functional preferences. Notably, valuation- and social-related motifs showed greater subdivision at higher K values, whereas the affective motif remained comparatively cohesive, consistent with its more spatially compact posterior distribution. In contrast, K = 3 provides the minimal resolution at which posterior affective, middle valuation, and anterior social motifs are simultaneously isolated as spatially coherent and functionally distinct subregions, consistent with quantitative model-selection criteria.

Together, these meta-analytic results provide convergent evidence for a tripartite motif-based functional organization of the VMPFC. This organization integrates affective processing, value computation, and social cognition along a continuous posterior–anterior axis, establishing a principled foundation for subsequent individual-level validation and connectivity-based analyses (Studies 2–4).

### Study 2 | Validation of the tripartite organization at the individual level

To confirm that the tripartite motif-based organization of the VMPFC extends beyond meta-analytic aggregation of task-evoked activations across thousands of published fMRI studies, we next examined individual-level task fMRI (tfMRI) data from the Human Connectome Project (i.e., HCP Young Adult dataset, n = 667). Three tfMRI paradigms were selected to probe distinct functional processes associated with the VMPFC: affective processing (HCP Emotion), value-based decision making (HCP Gambling), and social cognition (HCP Social)^30^. For visualization, group-level task activation maps revealed a consistent posterior–anterior ordering of peak responses within the VMPFC (Fig. 2a), mirroring motif-consistent functional biases: the posterior region co-activated with the amygdala (AMG), insula (INS), and anterior cingulate cortex (ACC) during affective processing; the middle region with the ventral striatum (VS) and orbitofrontal cortex (LOFC) during reward processing; and the anterior region with the dorsomedial prefrontal cortex (DMPFC), temporoparietal junction (TPJ), precuneus (PreC), and anterior temporal lobe (ATL) during social processing. At the single-subject level, VMPFC peak activations along the posterior–anterior axis exhibited a robust tri-modal spatial distribution (Fig. 2b). A k-nearest-neighbor (KNN) classifier—which assigns each subject’s VMPFC activation peak to one of three task-defined labels based on spatial proximity within a smoothed, population-derived VMPFC labeling field—reliably discriminated the three functional motifs, achieving classification accuracies of 0.688 (left VMPFC) and 0.716 (right VMPFC), far exceeding permutation baselines (∼0.33, p < 0.001; see *Methods: HCP task fMRI data and KNN classification*). Importantly, these findings were not specific to the KNN classifier. Repeating the analyses using Support Vector Classification and Logistic Regression yielded comparable decoding accuracies in both hemispheres (Supplementary Fig. 3b), demonstrating that individual-level validation of the tripartite motif structure is robust across classifier families. These results demonstrate that the tripartite VMPFC architecture is reproducible across individuals and hemispheres.

**Figure 2.**
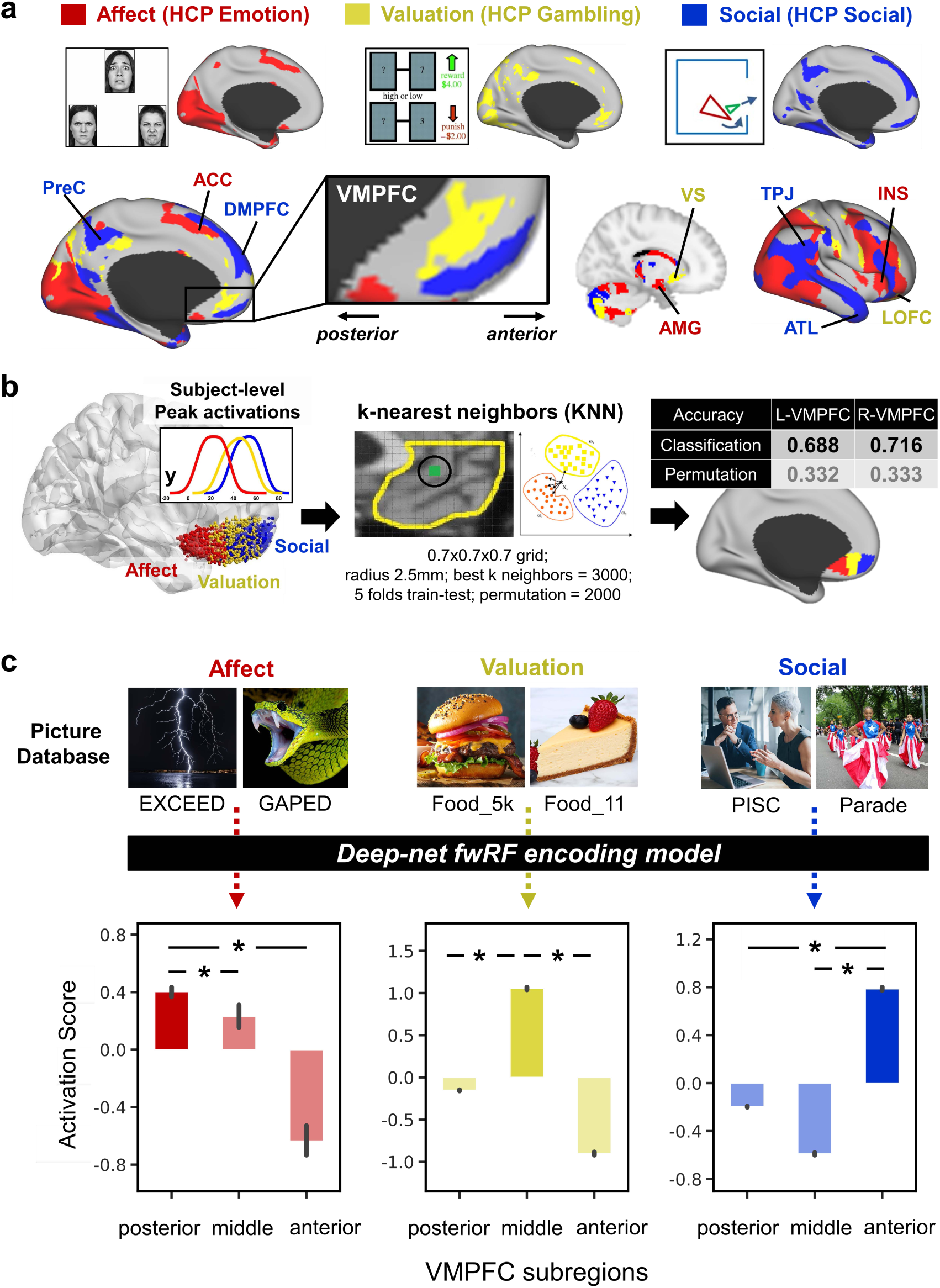
Individual-level validation and ANN-based encoding confirm a tripartite motif organization of the VMPFC in Study 2. **a,** Group-level standard activation maps for three HCP task fMRI paradigms show partially overlapping but dissociable functional systems corresponding to affective (red), valuation-related (yellow), and social (blue) processing. Canonical functional network nodes are annotated: emotion circuitry (INS, AMG, ACC), reward system (VS, LOFC), and social brain network (TPJ, ATL, DMPFC, PreC). Within the VMPFC (zoom-in view at the center), winner-take-all maps based on Cohen’s d visualize the dominant spatial weighting of each task domain, revealing a posterior-to-anterior ordering of motif dominance: posterior VMPFC is preferentially engaged by affective tasks, middle VMPFC by valuation tasks, and anterior VMPFC by social tasks. Standard activation and overlap maps are provided in Supplementary Fig. 2, confirming that these task activations are partially overlapping rather than anatomically segregated. **b,** Single-subject validation and classification. Subject-level peak activations were extracted for the three tasks, showing distinct distributions along the posterior–anterior axis (kernel density plotted over MNI y coordinate): posterior affect, middle valuation, and anterior social. A k-nearest-neighbors (KNN) classifier trained on within-subject activation features reliably assigns VMPFC motif labels in both hemispheres. **c,** ANN (deep-net fwRF) encoding. An image-driven encoding model (derived from the NSD database) predicts subregion-specific responses to classical, publicly available image databases dominated by emotional content (e.g., EXCEED, GAPED), value-related reward content (e.g., Food_5k, Food_11), or social content (e.g., PISC, WIDER-Parade). Predicted responses show motif-selective tuning to different stimulus domains, with posterior VMPFC responding most strongly to emotional stimuli, middle VMPFC to valuation stimuli, and anterior VMPFC to social stimuli. These results show that the tripartite VMPFC motif organization generalizes to ecologically rich, naturalistic stimuli. **Note: All face images are computer-generated using thispersondoesnotexist.com.**

Although task-based fMRI provides a powerful test of domain specificity, such paradigms necessarily sample a limited and experimenter-defined set of psychological conditions. To assess whether the tripartite motif-based organization of the VMPFC reflects a more general principle of neural representation—one that extends to continuous, high-dimensional stimulus spaces—we complemented task fMRI with encoding models trained on naturalistic stimuli. This approach allows us to ask whether VMPFC subregions exhibit systematic and predictable response preferences when driven by rich visual inputs that more closely approximate real-world experience. To operationalize this test, we leveraged artificial neural-network (ANN) encoding models trained on large-scale naturalistic stimuli (Fig. 2c). Building on recent evidence that deep networks approximate representational transformations in high-level cortex^31,32^, we trained feature-weighted receptive-field models (fwRF, see Extended Data Fig. 3a) on the Natural Scenes Dataset (NSD), which provides 22,000–30,000 stimulus–response pairs from 7T fMRI^33^ (see *Methods: ANN-based encoding model*). Separate models predicted the mean response of the posterior, middle, and anterior VMPFC subregions using hierarchical visual features extracted from AlexNet, spanning low-level image properties (e.g., edges and textures) to higher-level object- and scene-related representations. When applied to more than 30,000 naturalistic images drawn from publicly available emotional, value-related, and social picture databases (Supplementary Table 2), the predicted activation profiles—obtained by passing each image through the trained encoding model to estimate its expected fMRI response in each VMPFC subregion—reproduced the posterior→middle→anterior progression of functional dominance observed in HCP tasks: emotional images maximally engaged posterior VMPFC, value-related images engaged middle VMPFC, and social images selectively engaged anterior VMPFC (all *p*s < 0.001, see Figure 2c).

Because real-world stimuli rarely map cleanly onto a single domain, we next asked whether VMPFC subregions exhibit systematic combinatorial responses when multiple functional dimensions co-occur within a single image. We therefore tested motif-combinatorial selectivity by probing the trained encoding models with curated natural image sets that jointly contained two or more psychological attributes (Extended Data Fig. 3b). Social × value stimuli (e.g., attractive human faces) co-activated anterior and middle VMPFC; value × affect stimuli (e.g., antique/aesthetic objects) engaged middle and posterior VMPFC; social × affect stimuli (e.g., moral images) recruited anterior and posterior VMPFC; and images containing all three functional dimensions (e.g., erotic images) robustly activated all subregions. These ANN-derived response patterns demonstrate that the VMPFC’s tripartite motif-based architecture is not unique to HCP tasks but generalizes to ecologically rich visual inputs and exhibits differential, combinatorial responses when multiple psychological dimensions co-occur, consistent with an architecture optimized for integration rather than strict functional segregation.

Together, these results establish that the tripartite VMPFC architecture identified in Study 1 meta-analysis is replicable at the level of individual HCP subjects, predictable by ANN models, and systematically modulated by domain combinations, highlighting its integrative role in linking affective, value-based, and social computations.

### Study 3 | Connectivity-based mechanisms underlying VMPFC individual differences

Although Studies 1-2 established a consistent anterior-middle-posterior organization of the VMPFC at the population level, closer inspection revealed substantial inter-individual variability in the precise spatial extent and boundary transitions of these motifs across participants (Fig. 3a). Understanding the origin of this variability is essential for identifying the network-level principles that shape VMPFC functional specialization. A large body of prior work suggests that a region’s functional response properties are strongly constrained by its connectivity architecture^34–37^. We therefore tested whether voxel-wise functional selectivity within the VMPFC can be explained by each voxel’s whole-brain connectivity fingerprint.

**Figure 3.**
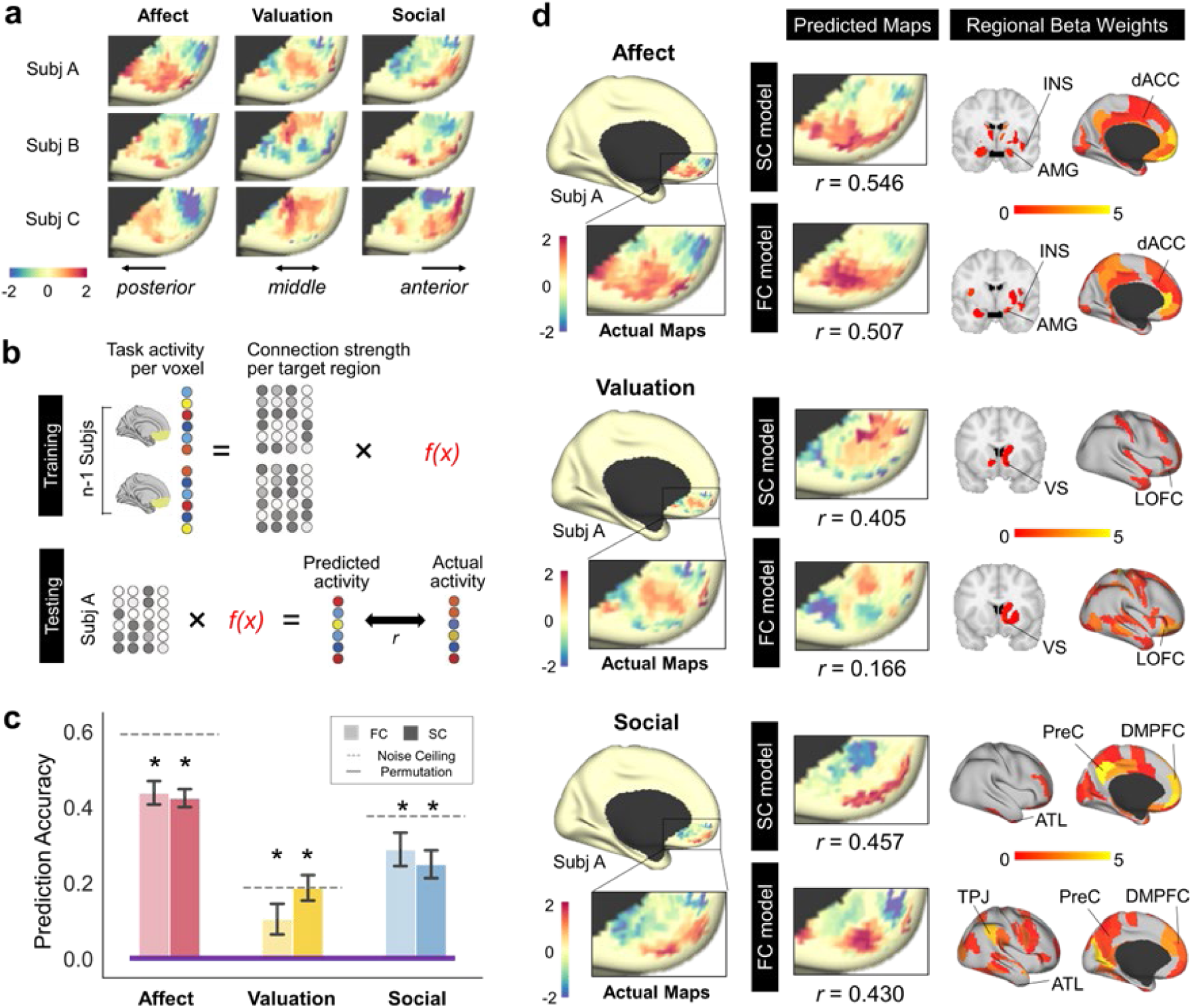
Connectivity fingerprints predict individual variation in VMPFC functional organization in Study 3. **a**, Task activation maps from representative HCP participants. A posterior–middle–anterior ordering of motif dominance is consistently observed across task domains (columns), alongside substantial inter-individual variability in activation topography (rows). Color bar represents task activation as GLM z-statistics. **b,** Voxelwise linear regression was used to predict VMPFC task activation from whole-brain connectivity profiles. During training, voxelwise task-activation values from n-1 participants (left; color-coded circles represent domain-specific task responses) were regressed onto corresponding connectivity fingerprints, defined as each voxel’s structural or functional connection strengths to all target brain regions (gray-scale circles), yielding a predictive model f(x). During testing, the learned model was applied to the left-out participant’s connectivity fingerprints to generate predicted voxelwise activity values. Model performance was quantified as the Pearson correlation (r) between predicted and observed activation maps within the VMPFC. This procedure was iterated across participants in a leave-one-subject-out cross-validation scheme. **c,** Model performance (r) for each task domain. Dashed lines denote the noise ceiling (run 1 × run 2 test–retest reliability for each HCP task); purple lines indicate permutation baselines. Both SC and FC models significantly outperform chance. **d,** Predicted maps and regional β-weights. Predicted activation maps from SC and FC models closely reproduce the observed VMPFC topography across task domains (color scale identical to **a**). Corresponding β-weight maps reveal distinct connectivity-defined functional motifs. Regional weights represent the absolute values of regression coefficients, converted to z-scores. The maps visualize the top 30 regions with the highest predictive contributions. Distinct motifs include: posterior VMPFC preferentially weighted by projections from the emotion circuitry (amygdala, insula, and dorsal ACC); middle VMPFC by the reward system (ventral striatum and lateral orbitofrontal cortex); and anterior VMPFC by the social brain network (precuneus, temporoparietal junction, anterior temporal lobe, and dorsomedial prefrontal cortex).

Because voxel-wise connectivity-based prediction modelling places more stringent demands on fMRI signal quality than localization or decoding analyses—particularly in ventral prefrontal and orbitofrontal cortex, where susceptibility artifacts and signal loss are prominent^38^—we collected a complementary in-house dataset (n = 42) using an optimized distortion-corrected acquisition^13^ and extended scanning duration^14^ to enhance data fidelity in the VMPFC (see Methods & Supplementary Methods). Importantly, this choice reflects the increased sensitivity of connectivity–activity mapping to signal-to-noise ratio and data density, rather than limitations of existing large-scale datasets. The optimized protocol substantially improved temporal signal-to-noise ratio (tSNR) in the VMPFC and yielded task activation patterns closely matching those observed in the HCP dataset (Extended Data Fig. 4). For each VMPFC voxel, we then computed structural connectivity (SC) fingerprints using diffusion tractography and functional connectivity (FC) fingerprints using resting-state fMRI, defined as voxel-wise connectivity strengths to all parcels in the brain.

To test whether connectivity architecture predicts functional responses, we used a voxel-wise least-squares linear regression framework^36^ to estimate task-evoked activation from connectivity fingerprint, applying leave-one-subject-out cross-validation (Fig. 3b). Both structural and functional connectivity profiles significantly predicted VMPFC activation patterns across all three tasks (all ps < 0.001; Fig. 3c), indicating a reliable relationship between connectivity architecture and functional response profiles. Importantly, the aim of this analysis was not to maximize predictive accuracy, but to preserve interpretability of regression coefficients so as to identify which large-scale networks preferentially contribute to each VMPFC motif.

Consistent with this goal, regression β-weight maps revealed highly structured and spatially specific connectivity–function relationships. Voxels in the anterior VMPFC were best predicted by connectivity with canonical social brain network regions, including dorsomedial prefrontal cortex (DMPFC), precuneus (PreC), temporoparietal junction (TPJ), and anterior temporal lobe (ATL). Voxels in the middle VMPFC were preferentially predicted by connectivity with reward-related regions, including ventral striatum (VS) and lateral orbitofrontal cortex (LOFC). Voxels in the posterior VMPFC were best predicted by connectivity with affective and interoceptive circuitry, including amygdala (AMG), insula (INS), and dorsal anterior cingulate cortex (dACC) (Fig. 3d). These modality-specific connectivity fingerprints closely mirror the tripartite motif-based organization identified in Studies 1-2, providing a mechanistic account of how distributed network inputs give rise to motif-consistent functional specialization within the VMPFC.

To directly compare the two connectivity modalities, we quantified the spatial correspondence between SC- and FC-based model predictions. Across participants, SC-and FC-predicted activation maps showed consistently positive spatial correlations within the VMPFC across all three functional domains (Supplementary Fig. 5a; all mean *rs* > 0.33, all *ps* < 0.001), indicating that the two modalities converge on similar large-scale functional topographies. In contrast, correspondence between the underlying regional β-weight profiles was weak (all *rs* < 0.11, all *ps* > 0.10; Supplementary Fig. 5b), suggesting that structural and functional connectivity rely on partially distinct connectivity features while nevertheless recovering similar motif-level organizations. This pattern indicates that similar functional topographies can emerge from different underlying connectivity predictors.

The connectivity–activity mapping was robust across multiple analytical choices. Similar results were obtained using different whole-brain atlases (e.g., BNA and HOA) (Extended Data Fig. 5a) and models trained to predict voxel-wise task activation from connectivity fingerprints in the in-house dataset accurately generalized to the independent HCP cohort, demonstrating out-of-sample generalization across datasets (Extended Data Fig. 5c). Likewise, connectivity-based parcellation of the VMPFC consistently reproduced the same anterior–middle–posterior organization across datasets and modalities, including HCP resting-state fMRI, HCP 7T movie fMRI, HCP diffusion MRI, and corresponding in-house resting-state and diffusion data (Extended Data Fig. 6). Together, these findings suggest that the tripartite motif-based organization of the VMPFC is not arbitrary or task-specific, but instead arises from systematic variation in the relative influence of three large-scale networks converging onto distinct zones of the VMPFC.

Taken together, these findings indicate that individual differences in VMPFC functional organization reflect systematic variation in whole-brain connectivity architecture. The anterior VMPFC is preferentially shaped by projections from the social brain network, the middle VMPFC by the reward system, and the posterior VMPFC by the emotion circuitry. This provides a mechanistic explanation for how large-scale network inputs give rise to stable yet individually variable motif-consistent patterns of functional selectivity within the human VMPFC.

### Study 4 | Generality across lifespan, species, and imaging modalities

Studies 1-3 established a robust tripartite motif-based organization of the VMPFC in adults based on task activation and brain connectivity. In Study 4, we asked whether this architecture generalizes across developmental stages, species, and neurobiological modalities. Specifically, we examined whether the tripartite pattern persists from early development to late adulthood, extends to non-human primates, and can also be derived from other imaging features such as gray-matter morphology, myelination, brain entropy, and neurotransmitter distributions.

Using resting-state data spanning the entire human lifespan (including BCP, HCP-Development, HCP-Young Adult, and HCP-Aging cohorts; age range: 0-100 yrs, total n = 2,293) together with rhesus macaques data (PRIME-DE-Macaque Oxford dataset, n=19), we performed task-free, connectivity-based parcellation of the VMPFC. Across all human cohorts and species, we identified a consistent posterior–middle–anterior tripartite subdivision (Fig. 4a). Quantitatively, although K = 2 and K = 3 yielded nearly identical mean cross-dataset Dice similarity (0.718 vs. 0.719; Extended Data Fig. 7), K = 3 exhibited substantially lower variability across cohort pairs (SD = 0.073 vs. 0.124), indicating more uniform agreement across developmental stages. The greater dispersion at K = 2 was driven primarily by reduced similarity involving the infant cohort. Inspection of the corresponding parcellation maps revealed that the infant cohort exhibited a predominantly dorsal–ventral organization at K = 2, whereas older cohorts showed a more anterior–posterior pattern. One possible explanation is that coarse parcellation solutions are particularly sensitive to developmental changes in large-scale connectivity architecture, consistent with evidence that long-range cortico-cortical connectivity matures more gradually than local and subcortical circuitry^39,40^. At K = 3, however, all cohorts—including infants—showed a similar posterior–middle–anterior organization and substantially more uniform cross-cohort agreement, supporting this tripartite solution as the most stable and developmentally generalizable description of VMPFC organization.

**Figure 4.**
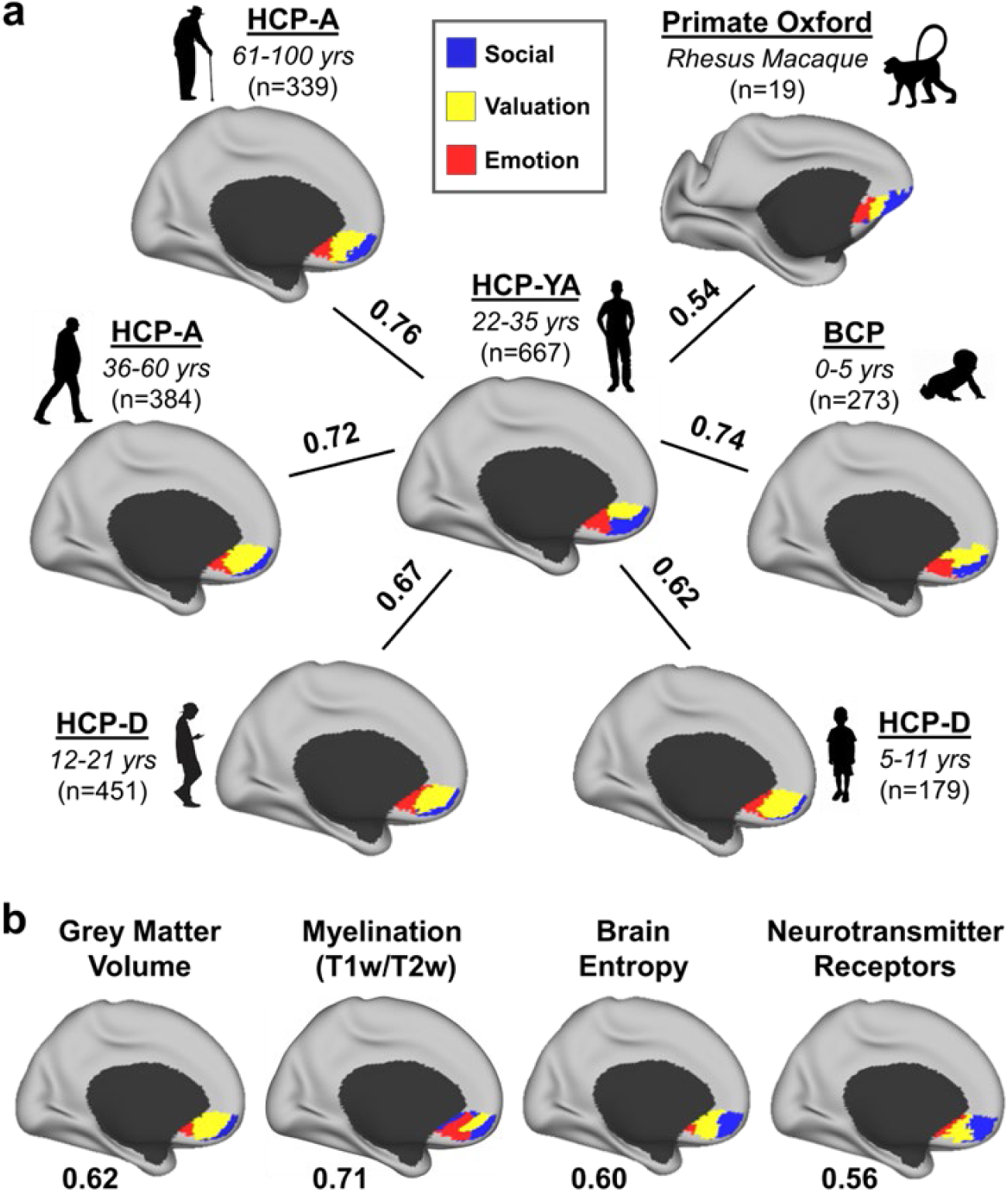
Lifespan, cross-species, and multimodal generality of the tripartite VMPFC motif organization in Study 4. **a**, Lifespan and cross-species parcellations. Task-free, connectivity-based parcellations of the VMPFC from resting-state fMRI reveal a reproducible three-parcel configuration across human development (i.e., HCP lifespan datasets) and in non-human primates (i.e., PRIME-DE dataset). Parcels follow a posterior–middle–anterior arrangement matching the young-adult reference (center). Solid lines, along with their adjacent values, indicate the maximum macro-averaged Dice similarity relative to the HCP-YA parcellation atlas. **b,** Parcellation based on multimodal brain features. Complementing the parcellations obtained in Studies 1–3 from task activation and connectivity, parcellations computed from other imaging features—gray-matter covariance, myelination-index covariance (T1w/T2w), temporal entropy of BOLD, and neurotransmitter receptor covariance—each recover a comparable posterior–middle–anterior tripartition of the VMPFC. The number underneath indicates Dice similarity relative to the HCP-YA parcellation atlas. This convergence indicates that the tripartite functional motif organization of the VMPFC is detectable across multiple neuroimaging dimensions.

We next tested whether the same organization could be recovered from diverse neurobiological features. Using the HCP Young Adult Multimodal Dataset and Hansen Receptors Atlas, we repeated the analysis with morphological (gray-matter volume covariance), microstructural (myelination index), dynamical (temporal entropy), chemoarchitectonic (neurotransmitter receptor density), and connectivity-based (functional and structural connectivity) measures (Fig. 4b). Internal clustering metrics favored different values of K across modalities (Supplementary Fig. 1), indicating that distinct features capture different spatial scales of VMPFC organization. Connectivity-based measures preferentially supported coarser partitions (K = 2), whereas myelination favored finer subdivisions (K = 5), with gray-matter volume and brain entropy favoring intermediate resolutions near K = 3. Rather than representing inconsistency, this pattern suggests that different modalities emphasize partially overlapping but non-identical aspects of VMPFC architecture. To integrate evidence across modalities, we performed a rank-based aggregation analysis using both the Silhouette coefficient and Davies–Bouldin index (Supplementary Fig. 1b). Across all six modalities, K = 3 achieved the best aggregate ranking under both metrics, indicating that it represents the most consistently supported solution across feature spaces. Moreover, K = 3 maximized cross-modality convergence (Extended Data Fig. 8), showed strong reproducibility across independent datasets and developmental cohorts (Extended Data Fig. 7), and corresponded closely to the affective, valuation, and social motifs identified in Studies 1–3.

Together, these findings demonstrate that the tripartite VMPFC organization generalizes across development, species, and neurobiological modalities, suggesting that it reflects a fundamental organizing principle rather than a modality- or dataset-specific phenomenon. Given known differences in cortical expansion, cytoarchitecture, and areal boundaries between humans and non-human primates, these results should not be interpreted as evidence for strict homology of cortical fields, but rather for convergence in large-scale organizational principles of the VMPFC.

## Discussion

The ventromedial prefrontal cortex (VMPFC) has long been implicated in affect, decision making, and social cognition, yet a principled account of how these diverse functions are organized within a single cortical territory has remained elusive. Although previous studies have reported multiple VMPFC subdivisions (Extended Data Fig. 1), most have remained largely descriptive, identifying regional boundaries within a particular dataset or modality without establishing whether those boundaries reflect a general organizational principle that reproduces across individuals, measurement modalities, developmental stages, and species. Consequently, it has remained unclear whether the functional heterogeneity of the VMPFC reflects a collection of partially independent processes or a coherent underlying architecture. Importantly, the present work addresses a different level of organization from prior studies that focused on ventral–dorsal dissociations across the broader medial prefrontal cortex. While ventral and dorsal medial prefrontal systems are known to differ in their connectivity and functional profile ^41^, the unresolved question addressed here is whether the VMPFC itself contains an internal organizational structure capable of explaining its heterogeneous involvement across affective, valuation-related, and social functions.

The present findings suggest that this heterogeneity is organized by a reproducible anterior–middle–posterior motif architecture that generalizes across individuals, modalities, developmental stages, and species. Accordingly, the primary contribution of the present study is not the identification of three parcels per se, but the demonstration that a common tripartite motif architecture provides a reproducible organizational scaffold linking functional specialization, connectivity architecture, development, and multimodal neurobiology within the VMPFC. Several observations support this conclusion. First, using cross-validated connectivity-to-activity mapping, we show that voxel-wise functional topography within the VMPFC can be predicted from whole-brain connectivity fingerprints in individual participants, indicating that the tripartite organization emerges systematically from large-scale network architecture rather than from group-level averaging. Second, the same organizational structure converges across multiple independent modalities—including meta-analytic co-activation, task fMRI, ANN-based encoding models, structural and functional connectivity, and chemoarchitectonic features—demonstrating that the motifs are not specific to any single measurement framework. Third, the ANN-based naturalistic analyses show that these motifs generalize beyond constrained laboratory paradigms and exhibit systematic combinatorial recruitment when affective, value-related, and social information co-occur within the same stimuli. Together, these findings provide a unifying account of the diverse functions historically attributed to the VMPFC, suggesting that affective, motivational, and social processes reflect coordinated variation within a common large-scale architecture rather than a collection of unrelated psychological functions. More broadly, the tripartite motif framework offers a biologically grounded organizational scaffold for understanding VMPFC function across cognitive, developmental, and comparative levels of analysis.

The tripartite organization is supported by longstanding theoretical proposals and cross-modal empirical convergence—two complementary criteria for a biologically meaningful parcellation. Narrative syntheses have repeatedly suggested that the VMPFC contains separable circuits for affect regulation, valuation, and social cognition^1,6,21^, but these accounts largely integrated heterogeneous findings without formally testing spatial arrangements and boundaries, reproducibility, or biological grounding. Our meta-analytic mapping quantitatively formalizes this hypothesis by demonstrating that three motifs arranged along the anterior–posterior axis show preferential associations with affect-, value-, and social-related processes (Fig. 1). Importantly, while prior whole-brain and medial prefrontal atlases consistently resolve a three-part VMPFC topology (Extended Data Fig. 1), they do not establish functional specialization; here, specialization is captured by the domain-dominant decoding profiles and high interpretability of the three-motif solution. Notably, this tripartite structure is preserved across parcellation resolutions (Extended Data Fig. 2): posterior parcels consistently exhibit affect-related dominance, middle parcels valuation-related dominance, and anterior parcels social-related dominance, with lower K values merging motifs and higher K values subdividing them. In this context, K = 3 represents the minimal resolution that simultaneously isolates all three motifs as spatially coherent and functionally distinct subregions. Across lifespan cohorts and non-human primates, the same threefold layout shows the strongest cross-dataset correspondence among candidate parcellations (Extended Data Fig. 7), and multimodal features—including gray-matter covariance, myelination, temporal entropy, structural connectivity, and neurotransmitter receptor distributions—converge on a similar configuration (Extended Data Fig. 8). Although internal clustering indices vary across modalities (Supplementary Fig. 1), these metrics primarily reflect geometric properties rather than biological validity; accordingly, parcellation quality is best judged by reproducibility, interpretability, and convergence across independent markers^42,43^. Together, these findings support K = 3 as the most parsimonious and biologically grounded description of VMPFC organization.

Although a tripartite motif scaffold recurs across datasets and modalities, the precise spatial boundaries between motifs are not expected to align perfectly, and this variability is informative rather than a limitation of the framework. Different modalities emphasize distinct biological signals and operate at different spatial scales, such that each captures the same anterior–middle–posterior organization with slightly shifted boundary placements. At the individual level, our connectivity-activity analyses in Study 3 suggest a mechanistic source of this heterogeneity: variation in the relative strength of social-, reward-, and affect-related network inputs shifts the transition zones between motifs, preserving the overall tripartite layout while producing person-specific boundary locations. From a developmental and comparative perspective, partial (rather than perfect) alignment is also expected, given ongoing network differentiation across childhood and additional uncertainty in cross-species correspondence. These observations indicate that the tripartite motifs should not be interpreted as rigid functional modules, but as dominant organizing axes defined by systematic variation in the relative influence of large-scale networks along the anterior–posterior extent of the VMPFC. Future work that explicitly models boundary uncertainty and individual variability—using gradient-based^44^ or probabilistic approaches^45^—will be essential for capturing how this organizational scaffold flexibly adapts across individuals, development, and species.

At a mechanistic level, our results bridge local and global theories of VMPFC function. Traditional accounts have emphasized cytoarchitectonic heterogeneity (e.g., agranular, dysgranular, granular), whereas our connectivity-based analyses highlight how macroscale wiring shapes local specialization, in line with evidence that connectivity profiles can predict later functional selectivity in associative cortex^34–36,46^. In Marr’s terms, our tripartite map primarily addresses the *implementational* level, specifying where different classes of information preferentially enter the VMPFC: anterior regions receive stronger input from social and default-mode systems, middle regions from reward-related circuitry, and posterior regions from affective and interoceptive circuitry. At the *algorithmic* level, these inputs may be transformed via partially distinct but interacting computations. Posterior VMPFC may preferentially support reinforcement-learning–like prediction-error updating, affective appraisal, and visceral-state mapping ^5,47^. Middle VMPFC may integrate dynamic value and contextual information through cost–benefit appraisal mechanisms that transform multidimensional evidence into subjective utility ^16^. Anterior VMPFC may support mental simulation and relational inference, constructing abstract map-like representations of self, others, goals, and future possibilities that enable flexible model-based reasoning^16,48^. Importantly, the label “social” should not be interpreted as implying that the anterior motif is exclusively dedicated to social cognition. Rather, social and self-referential processes represent the dominant functional associations recovered by our meta-analytic and connectivity analyses. More generally, the anterior motif may participate in a broader class of computations involving latent-state inference, relational abstraction, cognitive-map construction, and mental simulation. This interpretation is consistent with recent computational accounts arguing that many regions traditionally assigned to the “social brain” are better characterized by the computations they perform than by the social nature of their inputs alone^49^. From this perspective, social cognition may represent one particularly important manifestation of a more general capacity for modeling agents, relationships, goals, and hypothetical future states. Together, these processes implement a broader computational goal of evaluating the self in context, consistent with “self-in-context” and “cognitive map” theories of medial prefrontal function^5,50^. This view is also compatible with the classic somatic marker hypothesis^2,51^ which proposed that VMPFC integrates visceral and affective signals to guide behavior under uncertainty. Our findings extend this framework by suggesting that emotional, motivational, and relational information converge through partially distinct but interacting motifs within the VMPFC. These computational interpretations remain provisional and will require future formal model-based tests. Future causal and cross-scale work—including lesion mapping, intracranial recording, and cross-species tract-tracing—will be crucial for evaluating these mechanistic hypotheses and linking VMPFC microcircuitry to its macroscopic functional organization.

The anterior–posterior organization identified here is also broadly consistent with established cytoarchitectonic gradients within primate medial prefrontal cortex^52,53^. A parsimonious interpretation of the present motifs is that anterior parcels overlap more strongly with frontopolar/BA10-like territories, whereas posterior parcels overlap more strongly with agranular and dysgranular cingulate-related cortex. Thus, the functional motifs identified here may partly reflect known anatomical differentiation along the medial frontal wall. Developmental and evolutionary considerations may provide a broader context for understanding the origins of these anatomical gradients. Posterior medial prefrontal territories are evolutionarily older and less differentiated, whereas anterior granular association cortex has undergone disproportionate expansion during primate evolution and exhibits prolonged developmental maturation^54^. Related proposals, including the dual-origin hypothesis of cortical development, suggest that large-scale cortical differentiation follows broad paleocortical and archicortical gradients^55–57^. Although the present data do not directly test these developmental or evolutionary mechanisms, they suggest that the tripartite motif organization may be embedded within broader anatomical gradients that shape the structure and function of the VMPFC.

This tripartite framework also helps reconcile the diverse psychological and physiological functions historically ascribed to VMPFC, including episodic memory, default-mode-network (DMN) engagement, interoception, and visceromotor control. Although VMPFC has often been linked to “memory,” meta-analytic and lesion evidence^58–60^ indicates that its role in autobiographical retrieval primarily reflects self-referential and value-based reconstruction rather than mnemonic storage mechanisms per se, which are more strongly associated with medial temporal lobe structures and adjacent, more dorsal medial frontal regions supporting map-like or mnemonic representations^61^. In our model, the anterior social subregion, specialized for self/other inference, and the middle valuation subregion, specialized for subjective worth and motivational significance, jointly support the simulation and evaluation of personally meaningful events, thereby bridging memory, affect and social cognition. Consistent with this view, ANN-based encoding showed that when the VMPFC was driven by images combining multiple functional dimensions, activity spread systematically across subregions: social × value stimuli preferentially engaged anterior + middle VMPFC, value × affect stimuli engaged middle + posterior VMPFC, social × affect stimuli engaged anterior + posterior VMPFC, and stimuli containing all three dimensions activated all parcels (Extended Data Fig. 3b). These motif-combinatorial responses indicate that neighboring subregions interact to integrate affective, value-related, and social information rather than operate as isolated modules. Importantly, the present framework does not propose that affective, valuation, and social processes are strictly separable or process-pure categories. These dimensions frequently overlap in natural cognition, and the tripartite motifs should instead be interpreted as dominant organizational tendencies within a highly interactive architecture. This interpretation is supported by standard activation and overlap maps from the HCP tasks, which reveal substantial overlap among affective, valuation-related, and social activations despite a consistent posterior–anterior bias in dominant VMPFC recruitment (Supplementary Fig. 2). It is further supported by the ANN-based naturalistic analyses, which demonstrate systematic combinatorial recruitment across motifs when multiple functional dimensions co-occur within the same stimulus (Extended Data Fig. 3b). From this perspective, partial overlap and combinatorial recruitment across motifs are expected properties of VMPFC organization rather than violations of the model. The same architecture naturally explains the VMPFC’s anchoring role within the DMN, where anterior sectors support self-projection and mentalizing, and middle/posterior zones supply evaluative and affective context for internally oriented cognition^1,58^. Meanwhile, the posterior affective subregion, densely connected with amygdala, hypothalamus and brainstem nuclei, maps visceral states and regulates autonomic activity, accounting for classical links to interoceptive awareness and bodily arousal^62,63^. Rather than reflecting unrelated domains, these mnemonic, introspective and autonomic phenomena thus emerge from coordinated interactions among three VMPFC streams, positioning the region as a hub where social relevance, motivational salience, and bodily feeling are integrated^5^.

The tripartite map further provides a useful scaffold for clinical translation. To explore its potential clinical relevance, we performed an exploratory mapping of independent clinical findings onto the VMPFC tripartite organization (Extended Data Fig. 10). Disease-related biomarkers and therapeutic targets associated with major depressive disorder, substance use disorder, and autism spectrum disorder exhibited spatial distributions broadly consistent with the affective, valuation-related, and social sectors of the VMPFC, respectively. In contrast, conduct disorder—a heterogeneous condition characterized by impairments spanning affective regulation, reward processing, and social cognition—showed findings distributed across all three sectors, with symptom dimensions preferentially localizing to different motifs. Although preliminary and based on a limited number of studies, these observations suggest that distinct psychopathological dimensions may preferentially engage different VMPFC motifs and provide initial support for the clinical relevance of the proposed framework.

Explicitly considering this subregional organization may improve the specificity with which clinical phenotypes are mapped onto underlying neural circuits, paralleling recent efforts to distinguish dorsal and ventral subdivisions of the posterior cingulate cortex in psychiatric research^64^. For example, neuromodulatory interventions, including deep brain stimulation and transcranial magnetic stimulation, may benefit from targeting subregion-specific connectivity patterns rather than treating the VMPFC as a unitary structure, and pretreatment activation or connectivity profiles within these motifs may help predict differential responses across pharmacological and psychotherapeutic interventions^6,8^. At the individual level, precision-fMRI approaches that derive stable, person-specific parcellations^65^ may enable reproducible biomarkers for affective, motivational, and social symptom dimensions. These translational implications should be viewed as hypotheses for future empirical testing rather than as immediate clinical applications, but they illustrate how a subregion-resolved map of the VMPFC can guide mechanistically grounded, transdiagnostic research in mental health.

Several limitations should be noted. First, the term “VMPFC” does not refer to a single cytoarchitectonic field but spans multiple partially overlapping areas (BAs 10, 11, 12, 14, 25, 32) with substantial interindividual variability in sulcal morphology^7,10,66^. Following prior consensus recommendations^1^, we defined a mask corresponding to the ventromedial sector most consistently sampled in functional imaging and meta-analytic databases, but future work using individualized, morphology-based delineations will be needed to refine the exact boundaries of the tripartite map^42^. Second, the gambling task paradigm used to probe valuation in Study 3 showed only modest test-retest reliability at the individual level (as reflected by the noise ceilings in Extended Data Fig. 5), limiting power for fine-grained individual-difference analyses. Future studies could enhance reliability by combining multiple value-based tasks within the same individuals and by adopting precision-fMRI designs that use dense, repeated sampling to derive stable, person-specific connectomic fingerprints of VMPFC function^65^. Third, although the psychological constructs used to interpret the motifs—affect, valuation, and social cognition—are supported by data-driven clustering of meta-analytic terms in both semantic and neural representational spaces (Extended Data Fig. 9), they remain broad descriptive categories rather than process-pure computational variables. Affective motifs may be more precisely modeled using valence, arousal, salience, interoceptive state, or appraisal variables; valuation motifs using subjective value, expected utility, reward prediction error, or cost–benefit integration; and social motifs using latent variables related to self–other inference, mental-state attribution, relational abstraction, or social prediction. Replacing broad domain labels with such model-derived variables would move the present framework from functional description toward a more explicit computational account of VMPFC organization^67,68^. Fourth, the reverse-inference analyses are inherently constrained by the composition of the existing neuroimaging literature. Because databases such as Neurosynth, BrainMap, and NeuroQuery are constructed from published studies, functions that have been less frequently investigated in task-based fMRI may be underrepresented in term-based rankings. Consequently, some VMPFC functions—particularly those that are rarely studied or reported—may be underestimated by the present meta-analytic framework. In addition, our ranking procedure was based on the mean posterior probability across all VMPFC voxels. While this provides an interpretable summary of the dominant functional motifs represented within the VMPFC, it is inherently more sensitive to terms associated with spatially extensive activation patterns than to terms linked to highly localized subregions, whose contributions may be diluted by averaging across the entire region. Accordingly, the decoded term rankings should be interpreted as reflecting dominant region-level associations rather than an exhaustive characterization of all functions represented within the VMPFC. Finally, although the tripartite pattern generalized across lifespan cohorts and non-human primates, its developmental and state-dependent trajectories remain unknown. Longitudinal imaging combined with naturalistic paradigms and computational modeling will be essential for tracing how posterior (affect), middle (valuation) and anterior (social) streams differentiate and reintegrate over time. Ultimately, the value of this architecture will be determined by whether it can not only organize prior findings but also prospectively predict behavior, clinical symptom profiles and treatment response across individuals.

## Methods

### Definition of the VMPFC mask

To ensure anatomical precision and comparability across studies, we adopted a VMPFC mask defined by a collection of experts as part of their Nature Neuroscience viewpoints article on the human VMPFC^1^. This mask was based on the Harvard–Oxford atlas, encompassing the frontal pole, medial and orbital frontal cortex, anterior cingulate, paracingulate, and subcallosal cortex, while excluding voxels dorsal to z = 0 mm or lateral to |x| > 12 mm. This definition has been widely adopted in meta-analytic work. Alternative VMPFC masks used in the literature^45,59,69,70^ produce similar coverage but differ in spatial extent; the present choice ensures maximal overlap with established meta-analytic conventions and avoids inclusion of dorso-medial or orbito-lateral territories.

The present study focused specifically on the VMPFC rather than the entire medial prefrontal cortex (MPFC) because prior work has already established robust functional and connectivity dissociations between ventral and dorsal medial prefrontal systems. Accordingly, the primary unresolved question motivating the present work was not whether ventral and dorsal MPFC differ, but whether the VMPFC itself contains a reproducible internal organization capable of accounting for its heterogeneous involvement across affective, valuation-related, and social functions. Our ROI definition was therefore hypothesis-driven and designed to isolate the ventromedial territory most consistently implicated across these functions while avoiding extension into broader dorsomedial systems that operate at a different anatomical and functional scale.

### Reverse-inference mapping of functional terms

Analyses were performed using the Neurosynth database (www.neurosynth.org) containing 14,371 fMRI studies automatically annotated with activation coordinates and psychological terms^71^. We employed the NS+ toolkit (https://github.com/meng-du/NSplus) for large-scale meta-analytic computation, allowing reproducible reverse-inference and co-activation modeling directly within the Neurosynth framework^59^.

To identify functions most consistently associated with the VMPFC, we computed reverse-inference maps for all 1,335 psychological terms in the database. For each voxel, posterior probabilities P (term | activation) were calculated assuming a uniform prior of 0.5, following the NS+ implementation of Bayesian inference. Terms were ranked by mean posterior probability across all voxels in the VMPFC mask, excluding anatomical or ambiguous non-function labels (e.g., orbital, limbic, mode, connectivity, state, trait). The top 15 terms (Fig. 1a) and top 50 terms (see Supplementary Table 1) were categorized into the social, valuation, and affective categories based on semantic similarity and prior usage in the literature, and visualized as probability maps. This provided an unbiased, data-driven summary of the region’s dominant psychological associations.

### Data-driven clustering and labeling of functional terms

To validate the functional categorization of Neurosynth terms, we performed three complementary analyses (Extended Data Fig. 9). First, each term was embedded in semantic space using Qwen3-Embedding-8B^72^. To reduce ambiguity from everyday word meanings, terms were embedded within the fixed template “cognitive neuroscience concept of {term}”. Hierarchical agglomerative clustering was then applied to the resulting embedding vectors using cosine distance, without imposing any predefined functional labels.

Second, we performed an analogous clustering analysis using whole-brain Neurosynth association maps. For each term, we obtained its term-based association map and represented the term as a vector of voxel-wise association values. To avoid circularity with the VMPFC parcellation, voxels within the VMPFC mask were excluded before clustering. Hierarchical agglomerative clustering was then applied using correlation distance, again without imposing any predefined functional labels.

Third, to reduce investigator bias in naming the recovered clusters, we conducted an automated labeling procedure using an independent large language model (Gemini 3 Pro). Clusters were anonymized as “Cluster 1,” “Cluster 2,” and “Cluster 3,” and the model was provided only with the constituent terms. It was instructed to assign each cluster a single broad functional label, excluding anatomical network names or narrow task-specific descriptors. The procedure was repeated 100 times for both the 15-term and 50-term solutions, and label consistency was summarized across runs.

### Parcellation based on Meta-analytic co-activation modeling (MACM)

Meta-analytic co-activation modeling (MACM) was used to quantify how frequently VMPFC voxels co-activate with other brain regions across diverse tasks ^27,70^. All analyses were performed in standard 2-mm MNI152 voxel space without using a predefined anatomical atlas, following established Neurosynth-based parcellation procedures ^73,74^.

The analysis was based on the full Neurosynth database (version 0.7; N = 14,371 studies). For each voxel in the brain, Neurosynth provides a study-wise activation profile indicating the studies in which that voxel was reported as active. To reduce noise and improve reliability, voxels associated with fewer than 150 studies were excluded. This procedure yielded 1,784 voxels within the VMPFC mask and 129,213 voxels outside the VMPFC.

For each VMPFC voxel, a whole-brain co-activation profile was computed by quantifying its co-activation relationship with every non-VMPFC voxel across studies. This produced a co-activation matrix of size 1,784 × 129,213, in which rows corresponded to VMPFC voxels and columns corresponded to non-VMPFC voxels. To reduce dimensionality and computational complexity, principal component analysis (PCA) was applied to this matrix and the first 100 principal components were retained. Each VMPFC voxel was therefore represented by a 100-dimensional co-activation feature vector summarizing its relationship with large-scale whole-brain activation patterns. The resulting dissimilarity matrix (1 – Pearson’s *r*) quantified voxel-wise differences between co-activation profiles.

Voxels were grouped using K-means clustering into solutions ranging from K = 2 to K = 6 parcels. To minimize sensitivity to initialization, clustering was performed with 1,000 random initializations and the solution with the lowest within-cluster variance was retained^73,75^. Optimal cluster number was evaluated using multiple validity metrics, including the Silhouette coefficient and Davies–Bouldin index^42,76^. The K = 3 solution simultaneously maximized cluster cohesion and separation, yielded spatially contiguous subregions arranged along the posterior–anterior axis (Fig. 1b), and showed the strongest convergence across independent modalities and datasets. Functional interpretation of the resulting parcels was subsequently examined using reverse-inference decoding.

### Functional decoding of VMPFC parcels

For MACM-derived VMPFC parcels, each parcel’s psychological profile was characterized by repeating the reverse-inference analysis within its submask. For each parcel, we extracted the highest-ranking functional terms and correlated its spatial map with Neurosynth term-based maps using Pearson correlation. The strength of each domain association was visualized as radar plots (Fig. 1d), allowing direct comparison across social, valuation, and affect axes. Across alternative K solutions (K = 2–6), we further generated word-cloud visualizations summarizing the top decoding terms per parcel. These analyses revealed that posterior parcels were consistently dominated by affect-related terms, middle parcels by valuation-related terms, and anterior parcels by social-related terms across resolutions. Lower-dimensional solutions (K = 2) merged these motifs into broader, mixed profiles, whereas higher-dimensional solutions (K ≥ 4) subdivided them into smaller parcels that retained domain preferences but showed increased fragmentation. In contrast, K = 3 provides the minimal resolution at which all three functional motifs are simultaneously isolated as spatially coherent and functionally distinct parcels, consistent with the tripartite organization of the VMPFC (Extended Data Fig. 2).

For resting-state FC (rsFC)-derived VMPFC parcels, we performed the same term-based decoding on parcel-specific resting-state connectivity maps. For each parcel and each participant, we extracted the parcel’s mean BOLD time series and computed voxel-wise Pearson correlations with all other brain voxels outside of VMPFC, yielding subject-level seed-based rsFC maps that were averaged to obtain a group-level map per parcel. Each group-level rsFC map was then decoded against the same set of 15 a priori social, valuation, and affect terms using the corresponding Neurosynth term-based maps. For each term, we correlated the parcel’s rsFC map with the term map and normalized parcel weights by dividing each weight by the sum across parcels, yielding proportional involvement scores that highlight the relative contribution of each parcel while controlling for overall term prevalence (Extended Data Fig 6d).

### HCP task fMRI data and KNN classification

Task-based analyses used the HCP Young Adult dataset^77,78^. Three paradigms capturing distinct VMPFC-relevant domains were analyzed: HCP Social (‘social – random’ contrast), HCP gambling (‘reward – punish’ contrast), HCP Emotion (‘emotion – baseline’ contrast). Group-level Cohen’s d effect-size maps were downloaded from the official S1200 Group Average Data Release. Group-level activation maps (Figure 2a) were visualized in Connectome Workbench and thresholded at d > 0.2 to delineate reliable activation.

To examine individual-level functional subdivisions, we extracted peak coordinates from subject-level contrast maps within the VMPFC mask (n=667 with significant VMPFC activations for all three tasks, see a similar method^79^). We implemented a prototype-based parcellation that assigns each voxel to one of three task-define labels (social, valuation, affect) according to spatially smoothed peak distributions. A uniform lattice of N = 54,072 grid points was created across the VMPFC. The objective was to estimate a label-probability matrix X (N × 3) satisfying spatial smoothness and data-fidelity constraints:

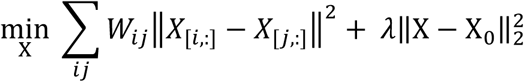

where W₍ᵢⱼ₎ = 1 if distance < 0.7 mm, 0 otherwise; X₀ is a local prior computed from activation peaks within r = 2.5 mm. The closed-form solution X⁎ = λ (L + λ I)⁻¹ X₀ uses the graph Laplacian L = diag(W 1) – W. λ was exponentially varied between 10⁻⁴ and 10⁻¹ to balance smoothness and fidelity.

Model performance was assessed by five-fold cross-validation. For each fold, 80% of peak coordinates served as training data to compute X₀ and X⁎; the remaining 20% constituted the test set. A k-nearest-neighbor classifier (k = 3,000) trained on labeled grid points predicted task labels for unseen peaks. Accuracy significance was established via 2,000 label-permutation tests per fold.

### ANN-based encoding model

To evaluate ecological generalization, we first established deep-network feature–weighted receptive-field (fwRF) encoding models^32^ (see Extended Data Fig 3a) on the Natural Scenes Dataset (NSD), a 7T fMRI resource in which eight adults each viewed 9,000–10,000 unique natural-scene photographs across 30–40 sessions (22,000–30,000 total trials per subject)^33^.

For each VMPFC subregion (posterior, middle, anterior; masks derived from Study 1 MACM parcellation), we trained a separate fwRF model predicting subregion-averaged fMRI responses. Specifically, the model integrates (i) multi-layer feature maps from a pretrained AlexNet, (ii) a 2D Gaussian pooling field specifying spatial weighting over feature maps, and (iii) ridge-regularized linear weights learned to optimize prediction accuracy. We used AlexNet-derived hierarchical visual features because they have been extensively validated in prior neural encoding studies of high-level visual and associative cortex, and because they permit transparent separation of feature complexity across network layers. Importantly, our conclusions do not depend on the semantic specificity or biological plausibility of any particular model architecture, but on relative differences in encoding performance across VMPFC subregions when driven by the same stimulus space.

To validate the performance of ANN-based fwRF encoding models, we quantified out-of-sample prediction accuracy on the NSD dataset using cross-validated Pearson correlation. Encoding models in the VMPFC typically yield modest predictivity compared to sensory regions due to susceptibility artifacts and complex functional selectivity. Consequently, we adopted a two-tiered validation strategy as a sensitivity analysis, balancing signal fidelity with population representativeness. For the primary analysis, we adhered to rigorous inclusion criteria (r > 0.05) based on recent large-scale high-fidelity encoding studies^80,81^, isolating participants with the most robust neural responses (3 out of 8 NSD participants). Secondarily, to prevent the exclusion of physiologically relevant but attenuated signals, we performed an analysis using a liberal threshold (r > 0.01, 6 out of 8 NSD participants), specifically selected to recover valid but weaker signals.

We next applied each trained fwRF model to predict each subregion’s responses to ∼30,000 domain-specific natural images encompassing social, value-related, and emotional contents (see full descriptions of all picture databases in Supplementary Table 1). Predicted responses were z-scored within subject. Domain selectivity was quantified using repeated-measures ANOVAs across subregions, followed by FDR-corrected post hoc comparisons. Additional mixed-domain image sets were used to test combinatorial selectivity (Extended Data Fig 3b), applying identical preprocessing and statistical procedures. Note that the prominent IAPS database (International Affective Picture System) was not included in the ANN-based encoding analyses because these images were reserved for in-house fMRI experiments in Study 3; as expected, their functional selectivity patterns were confirmed independently (Extended Data Fig. 4b). Crucially, the posterior–middle–anterior functional organization was reliably reproduced under both criteria (r > 0.05 or r > 0.01), confirming that the tripartite motif organization is a stable feature of VMPFC architecture that persists across varying levels of predictive fidelity.

### In-House Dataset with Optimized fMRI Acquisition and Task Paradigms

Forty-two healthy young adults (26 female; age 18–30; mean age = 22.6 ± 2.8 years) were recruited from Beijing Normal University. All participants reported normal or corrected-to-normal vision and no history of neurological or psychiatric conditions. Written informed consent was obtained in accordance with institutional ethics approval (IRB_A_0024_2020001).

All MRI data were collected on a 3T Siemens MAGNETOM Prisma scanner equipped with a 64-channel head coil at the BNU Brain Imaging Center. Because the VMPFC lies adjacent to orbitofrontal sinuses and is highly susceptible to signal drop-out, we implemented an optimized EPI protocol (see Supplementary Methods for details) to mitigate susceptibility signal loss in VMPFC^13,82,83^. Specifically, we used a shorter TE (22 ms)^12,84^ and a tilt of 30 degree relative to the AC–PC axis to reduce susceptibility gradients^85,86^. In addition, to improve within-subject reliability and signal stability in ventral prefrontal regions, we adopted an elongated scanning strategy with substantially increased task duration^14^. Compared with the standard HCP task-fMRI protocol (∼20 minutes total across tasks), our in-house protocol approximately doubled total task scanning time to ∼45 minutes per participant. This protocol substantially increased temporal Signal-to-noise (SNR) in the VMPFC relative to standard HCP sequences and produced activation patterns closely matching those observed in the HCP-Young Adult dataset (Extended Data Fig 4).

In-house participants completed three fMRI tasks probing social, valuation, and emotion processing. The social cognition and valuation tasks followed HCP designs, but with extended trial counts and longer task blocks to enhance within-subject reliability and sensitivity for connectivity-based predictive modeling, which is known to be particularly sensitive to data quality in ventral prefrontal cortex^14^. Based on converging evidence that posterior VMPFC is preferentially engaged during affect regulation rather than passive affect recognition^1^, we replaced the HCP emotion matching task with an emotional reappraisal task in which participants up-regulated or down-regulated affective responses to negative images in IAPS (International Affective Picture System). First-level GLMs were estimated in FSL, and subject-level activation maps were registered to native T1 space for subsequent modeling.

### Connectivity Fingerprinting: Predicting Activity from Connectivity

Structural connectivity was estimated using voxelwise probabilistic tractography implemented in FSL PROBTRACKX2 (5,000 streamlines per seed voxel; step size = 0.5 mm; curvature threshold = 0.2; distance correction enabled). Each voxel within the VMPFC served as an independent seed, and connection probabilities were computed to a whole-brain set of cortical and subcortical targets defined using the Brainnetome atlas^87^ combined with an MDTB cerebellar parcellation^88^. For functional connectivity, Fisher z-transformed Pearson correlations were calculated between the resting-state time series of each VMPFC voxel and each target parcel, yielding a subject-specific voxelwise functional connectivity fingerprint.

To test whether distributed connectivity profiles predict local functional specialization, we applied a least-square linear regression framework identical to prior connectivity–function mapping studies^34–36^. Each voxel constituted an observation, with its whole-brain connectivity vector used as predictors and its voxelwise task activation value as the dependent variable (Fig. 3b). Models were trained using leave-one-subject-out cross-validation: connectivity and activation data from N–1 participants were concatenated to estimate regression coefficients, which were then applied to the held-out participant’s connectivity fingerprints to generate a full predicted VMPFC activation map. Prediction accuracy was quantified as Pearson correlation between predicted and observed voxelwise activation patterns

Statistical significance was assessed with 1,000 permutation tests per fold, generated by randomly shuffling connectivity–activation pairings across participants to form empirical null distributions. Network contributions were inferred from standardized β-weight maps averaged across folds. These analyses revealed that anterior VMPFC activity was primarily predicted by connections to canonical social brain regions (precuneus, TPJ, ATL, dorsomedial PFC), middle VMPFC by reward system nodes (ventral striatum and lateral OFC), and posterior VMPFC by emotion circuitry (amygdala, insula, dorsal ACC). Predictive patterns were robust to atlas selection and replicated in independent HCP samples (Extended Data Fig. 5).

### Connectivity-Based Parcellation

Connectivity-based parcellation of the VMPFC was performed using CBPtools^76^, which implements voxel-wise connectivity fingerprint estimation and clustering. For each participant, connectivity profiles (i.e., voxel-to-voxel connectivity data) were derived separately from resting-state fMRI and diffusion MRI tractography (see illustration in Extended Data Fig 6a). Voxel-wise similarity matrices were constructed, and parcellation was performed using k-means clustering with k = 3, consistent with the tripartite organization established in Study 1. Group-level parcellations were computed by relabeling cluster identities across subjects using minimum Hamming distance and taking the voxel-wise mode. This procedure was performed both in the in-house dataset and in HCP-YA (3T resting fMRI, 3T DWI, and 7T movie fMRI), yielding a highly consistent posterior–middle–anterior VMPFC subdivision (Extended Data Fig 6b & 6c).

### Lifespan & Species Datasets

We leveraged publicly available lifespan cohorts and a non-human primate dataset to test the generality of the tripartite VMPFC organization. The human sample (downloaded on February 2021) comprised four large-scale cohorts from the HCP lifespan project^89^: BCP (ages 0–5, n=273), HCP-D (ages 5–21; n=630), HCP-Young Adults (ages 21–35; n=667), and HCP-Aging (ages 36–100, n=723). All human data (n=2,293) were minimally preprocessed according to the standard HCP pipelines and already aligned to a common MNI space. Because dataset acquisition and quality control details are extensive, we refer readers to prior publications^90–92^ rather than duplicate them here. In addition, we included resting-state fMRI data from Macaque Oxford dataset (rhesus macaques, n = 19) in PRIMatE Data Exchange^93^ to assess evolutionary conservation. All details about each dataset can be found in Supplementary Methods.

### Multimodal feature–based parcellation

To test whether the tripartite map could be identified from imaging features other than functional or structural connectivity, we leveraged HCP Young Adult Multimodal Dataset and Hansen Receptors Atlas and computed voxel-wise covariance or feature-profile similarity within the VMPFC based on: (1) gray-matter volume covariance networks, (2) cortical myelination indices (T1w/T2w ratio), (3) temporal entropy of resting-state BOLD time series (a measure of dynamical complexity), and (4) neurotransmitter receptor/transport-er density covariance (compiled from PET and autoradiography templates). Each feature set was used to compute a group-level parcellation of the VMPFC. All details about each multimodal feature can be found in Supplementary Methods and all data-driven model-selection metrics (e.g., Silhouette coefficient and Davies–Bouldin index) were provided in Supplementary Figure 1.

### Parcellation and Similarity Computation

For each HCP cohort or imaging modality, we computed group-level parcellations of the VMPFC across cluster resolutions K = 2–6. While human parcellations were derived in standard MNI space, the macaque parcellation required a tailored anatomical definition. Specifically, we mapped the human VMPFC mask to the macaque volumetric space via a sequential transformation (human volume to human surface to macaque surface to macaque volume). This process utilized a joint-embedding alignment framework to account for cross-species topological discrepancies and evolutionary areal expansion^94^.

To quantify similarity across parcellations, we employed the maximum macro-averaged Dice coefficient (equivalent to macro-F1) across all possible one-to-one permutations of cluster labels. Macro-averaging treats each cluster equally and guards against bias from cluster-size differences. For cross-species comparisons, we evaluated similarity on the common macaque cortical surface. To achieve this, both the derived macaque volumetric parcellation and the human reference parcellation were projected onto the human surface. This similarity metric underpins the cross-dataset and cross-modality stability analyses shown in Extended Data Fig. 7 and Extended Data Fig. 8, respectively.

### Robustness analyses and cross-database validation

To evaluate the stability of the MACM-derived VMPFC parcellation, we performed a series of robustness analyses varying both voxel-selection criteria and clustering algorithms. K-means was selected as the primary clustering approach because it is widely used in connectivity-based and meta-analytic brain parcellation studies, facilitates comparison with prior MACM investigations, and provides a transparent baseline method for identifying large-scale motif structure in low-dimensional co-activation spaces.

First, we varied the minimum-study inclusion threshold used for voxel selection in the MACM analysis (≥100, ≥150, and ≥200 studies per voxel), yielding VMPFC masks containing 2,211, 1,784, and 1,403 voxels, respectively. For each threshold, K-means clustering (K = 3) was repeated 100 times using independent random seeds. Spatial similarity between solutions was quantified using Dice coefficients relative to the consensus tripartite solution.

To assess dependence on algorithm choice, we repeated the parcellation analyses using Spectral Clustering and Agglomerative Clustering. For each algorithm, clustering was repeated 100 times and spatial similarity to the original K-means solution was quantified using Dice coefficients. Representative maps corresponding to the lowest-similarity solutions were retained for visualization to provide a conservative estimate of robustness.

To determine whether the MACM-derived VMPFC parcellation generalized beyond the Neurosynth dataset and whether the observed tripartite organization could be influenced by biases in the composition of the literature, we repeated the entire parcellation pipeline using an independent meta-analytic database, BrainMap. In contrast to Neurosynth, which is automatically constructed from high-frequency terms extracted from published articles, BrainMap is manually curated and provides human expert annotations of experimental paradigms and behavioral domains. Coordinate data were retrieved using Sleuth and converted into the same 2-mm MNI152 voxel space used for the Neurosynth analyses. The BrainMap database contained 4,341 studies and 22,301 experiments. Following the Neurosynth-based MACM pipeline, each voxel was represented by a binary activation vector across experiments. Voxels associated with fewer than 120 experiments were excluded from both the VMPFC and the rest of the brain, leaving 1,902 VMPFC voxels and 138,035 non-VMPFC voxels for analysis. PCA-based dimensionality reduction, voxel-wise co-activation profiling, K-means clustering, and cluster-validity assessment were then performed using the identical procedures described above for Neurosynth. Spatial correspondence between the resulting BrainMap-derived and Neurosynth-derived parcellations was quantified using Dice coefficients.

To evaluate potential dataset-composition biases, we additionally quantified the distribution of studies across affective, valuation, and social domains in both databases. For Neurosynth, domain assignments were derived from the data-driven term-clustering analyses described above using both the top 15 and top 50 VMPFC-associated terms. For BrainMap, studies were categorized according to the manually curated behavioral-domain and paradigm classifications. Domain counts were calculated separately for the entire database and for studies contributing to the VMPFC meta-analysis. We further quantified the distribution of individual experimental paradigms within VMPFC-activating studies to assess whether any single task disproportionately contributed to the observed parcellation structure (Supplementary Fig. 4).

To determine whether the individual-level validation depended on classifier choice, the subject-level decoding analyses in Study 2 were repeated using three supervised learning algorithms: K-nearest neighbors (KNN), Support Vector Classification (SVC), and Logistic Regression. KNN was selected as the primary classifier because it makes minimal assumptions regarding decision boundaries and provides an interpretable test of whether the group-level motifs can be recovered in individual participants. Classification accuracy was evaluated separately for the left and right VMPFC and compared against null distributions generated through permutation testing. Results are summarized in Supplementary Figure 3.

## Data and Code availability

All data and analysis code in this project have been deposited on the GitHub (https://github.com/BNU-Wang-MSN-Lab/VMPFC).

## Acknowledgement

We thank Gaolang Gong, Yong Liu, Ruiwang Huang, Tengda Zhao, Robert S. Chavez, Dale T. Tovar, Yunman Xia, Wenli Tao for valuable discussions and comments. This work was supported by the National Natural Science Foundation of China (32422033, 32430041, 32595491, 32000782 to Y.W.), the National Science and Technology Innovation 2030 Major Program (2022ZD0211000, 2021ZD0200500 to Y.W.), the Fundamental Research Funds for the Central Universities.

## Extended Figures

**Extended Data Fig. 1.**
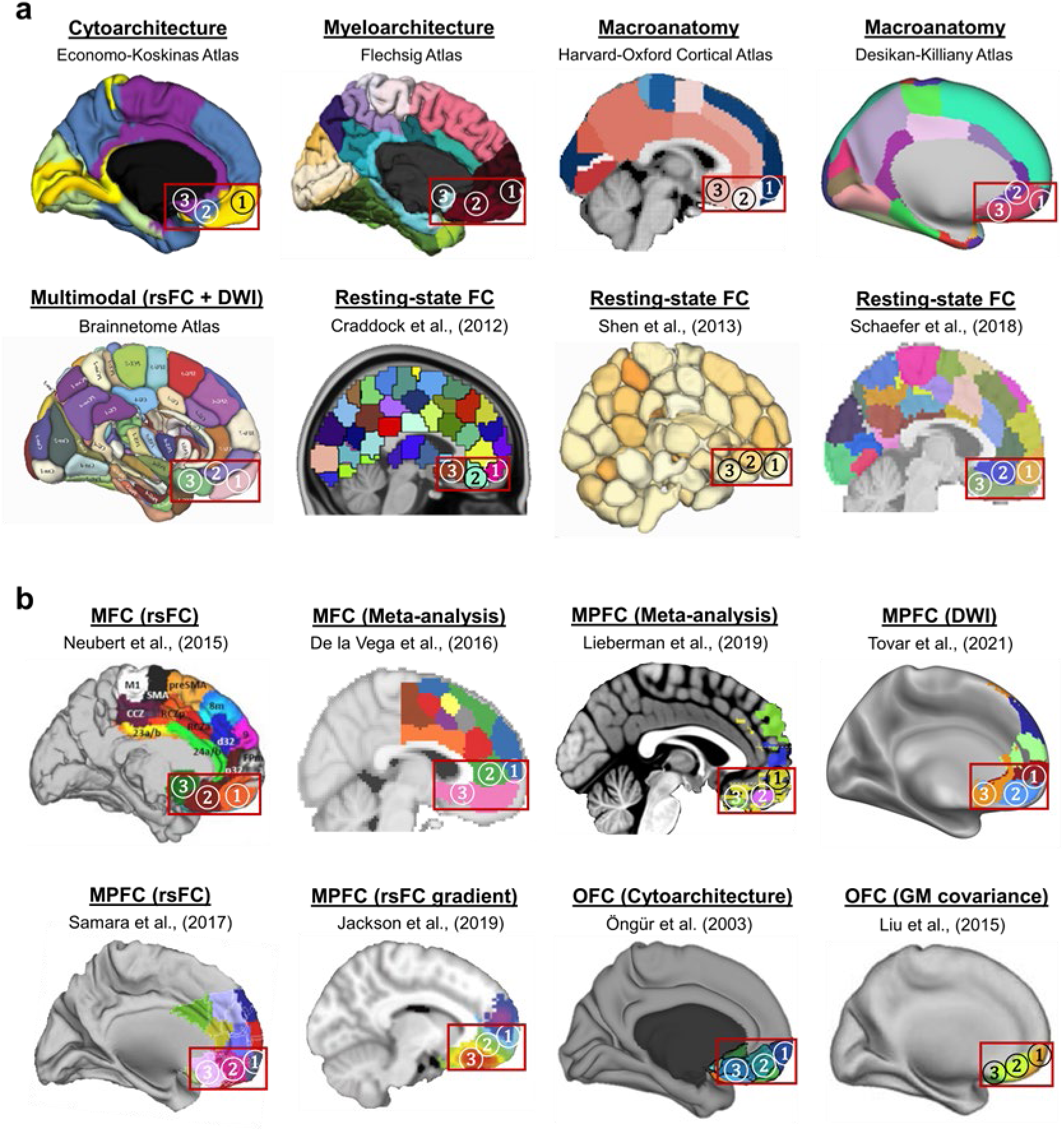
Converging evidence from diverse human atlases for a tripartite VMPFC. Panels compile representative **(a)** whole-brain and **(b)** regional medial/orbitofrontal parcellations derived from heterogeneous principles—including cyto-and myelo-architecture, diffusion tractography, resting-state functional connectivity, morphometric covariance, and gradient/meta-analytic methods. In each panel, the red bounding box marks the VMPFC; numerals 1-2-3 indicate the anterior, middle, and posterior sectors, respectively. Across atlases, the VMPFC reproducibly resolves into three contiguous parcels along the anterior-posterior axis, consistent with the tripartite organization established in Studies 1-4. See atlas references^9,45,59,74,95–101^

**Extended Data Fig. 2.**
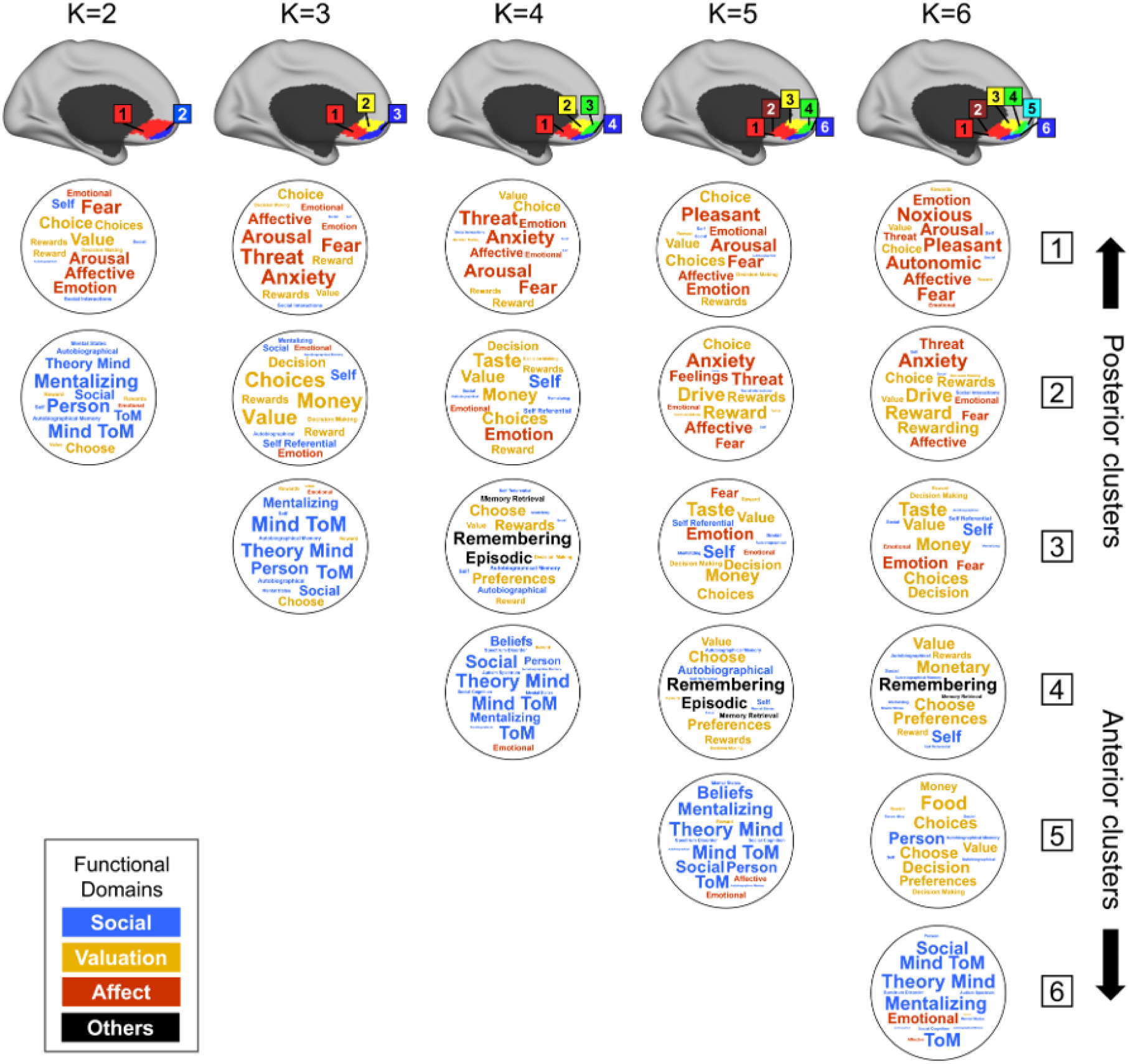
Functional decoding across parcellation solutions (K = 2–6) reveals stable motif structure with optimal interpretability at K = 3 in Study 1. Top: Schematic views of VMPFC parcels derived from MACM clustering for K = 2–6. Below each parcel, word clouds display the most diagnostic reverse-inference terms (font size reflects normalized decoding weight), color-coded by functional domain (blue = social, yellow = valuation, red = affect, black = others). Across all resolutions, posterior parcels are consistently dominated by affect-related terms, middle parcels by valuation-related terms, and anterior parcels by social-related terms, indicating that the three functional motifs are preserved across clustering solutions. Lower-dimensional solutions (K = 2) merge adjacent motifs into broader, mixed profiles, whereas higher-dimensional solutions (K = 4–6) subdivide them into smaller parcels that retain their dominant functional preferences. Notably, the affective motif remains comparatively cohesive across resolutions, whereas valuation and social motifs are progressively subdivided at higher K values, consistent with their broader spatial extent and greater internal heterogeneity within the VMPFC. In contrast, K = 3 provides the minimal resolution at which posterior affective, middle valuation, and anterior social motifs are simultaneously isolated as spatially coherent and functionally distinct subregions. This solution yields the clearest correspondence between functional decoding and spatial organization and converges with the quantitative model-selection results shown in Fig. 1b. Colors are consistent across figures; schematic views are shown in MNI orientation. For simplicity, results are displayed for the left VMPFC, which closely mirrors the right VMPFC.

**Extended Data Fig. 3.**
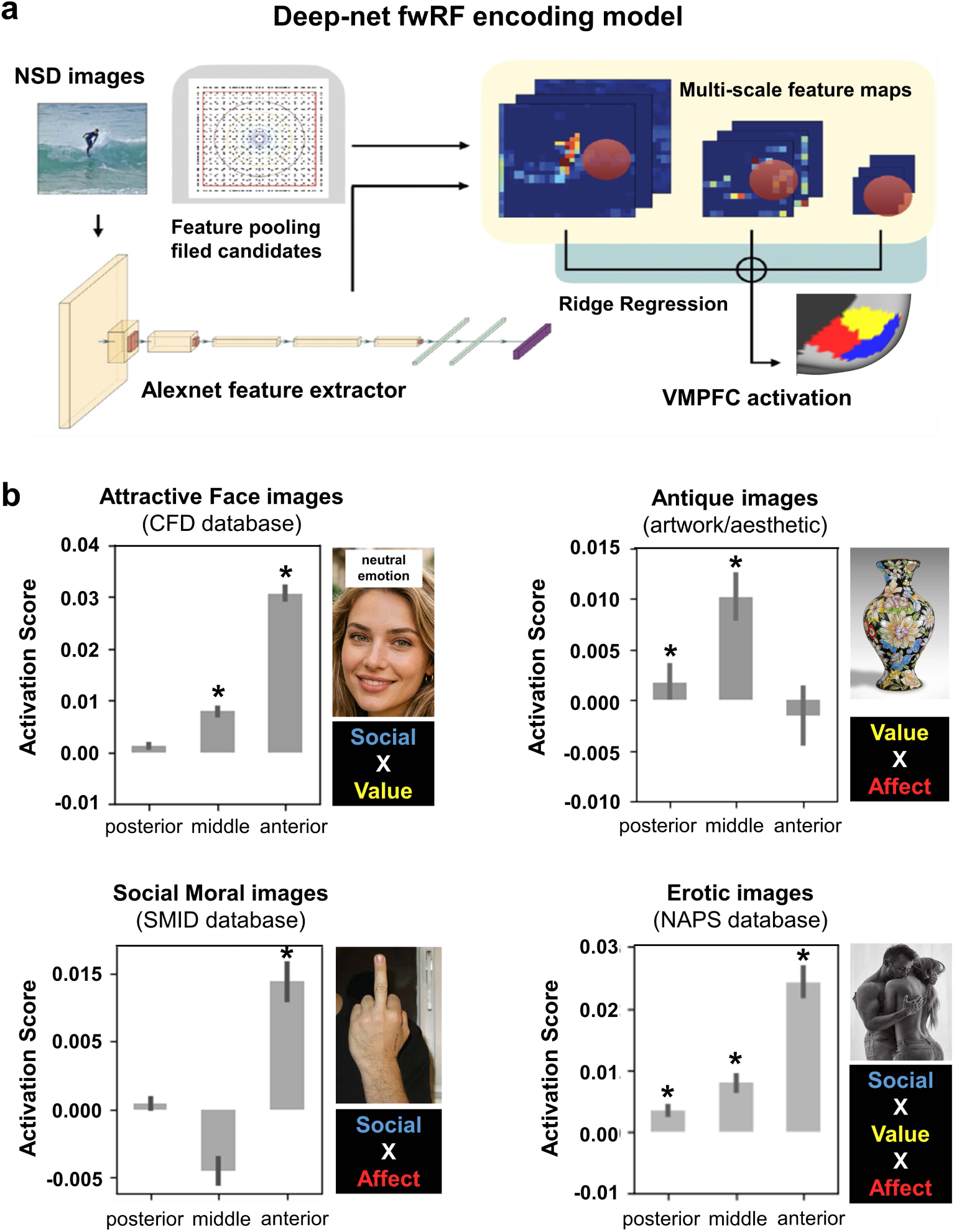
Deep-net fwRF encoding model and cross-dataset tests reveal motif-combinatorial responses in the VMPFC in Study 2. **a,** S Schematic of the encoding approach. Following established work^32^, we trained region-specific deep-net fwRF models to predict parcel-averaged fMRI responses in the posterior (pVMPFC), middle (mVMPFC), and anterior (aVMPFC) functional motifs. Natural scene images from NSD were passed through an AlexNet feature extractor to obtain multi-scale feature maps. Candidate 2-D Gaussian feature-pooling fields were combined with ridge regression to learn feature weights that best predict VMPFC activation; models were fit separately for each parcel and subject, following the fwRF procedure described in Methods. **b,** Probing the trained models with mixed-domain image sets. Bars show z-scored predicted responses (mean ± s.e.m.) for each VMPFC motif when the models are driven by images that combine two or three functional domains: attractive faces (CFD; neutral emotion selected), antique/artwork images, social moral images (SMID; negative valence), and erotic images (NAPS_ERO). Response profiles follow domain composition: attractive faces (Social × Value) preferentially engage anterior and middle VMPFC motifs with minimal posterior response; antique/artwork images (Value × Affect) engage middle and posterior motifs with minimal anterior response; social moral images (Social × Affect) engage anterior and posterior motifs; and erotic images (Affect × Value × Social) engage all three motifs. Asterisks indicate p < 0.05 significance above zero (post-hoc paired tests, FDR-corrected). Color labels denote domains (blue = social, yellow = valuation, red = affect); bars are gray for readability. Together, these results show that VMPFC motifs integrate information across domains in a differential, combinatorial manner, rather than responding as isolated, domain-specific modules, reinforcing the tripartite motif organization established in Fig. 2. **Note: All face images are computer-generated using thispersondoesnotexist.com.**

**Extended Data Fig. 4.**
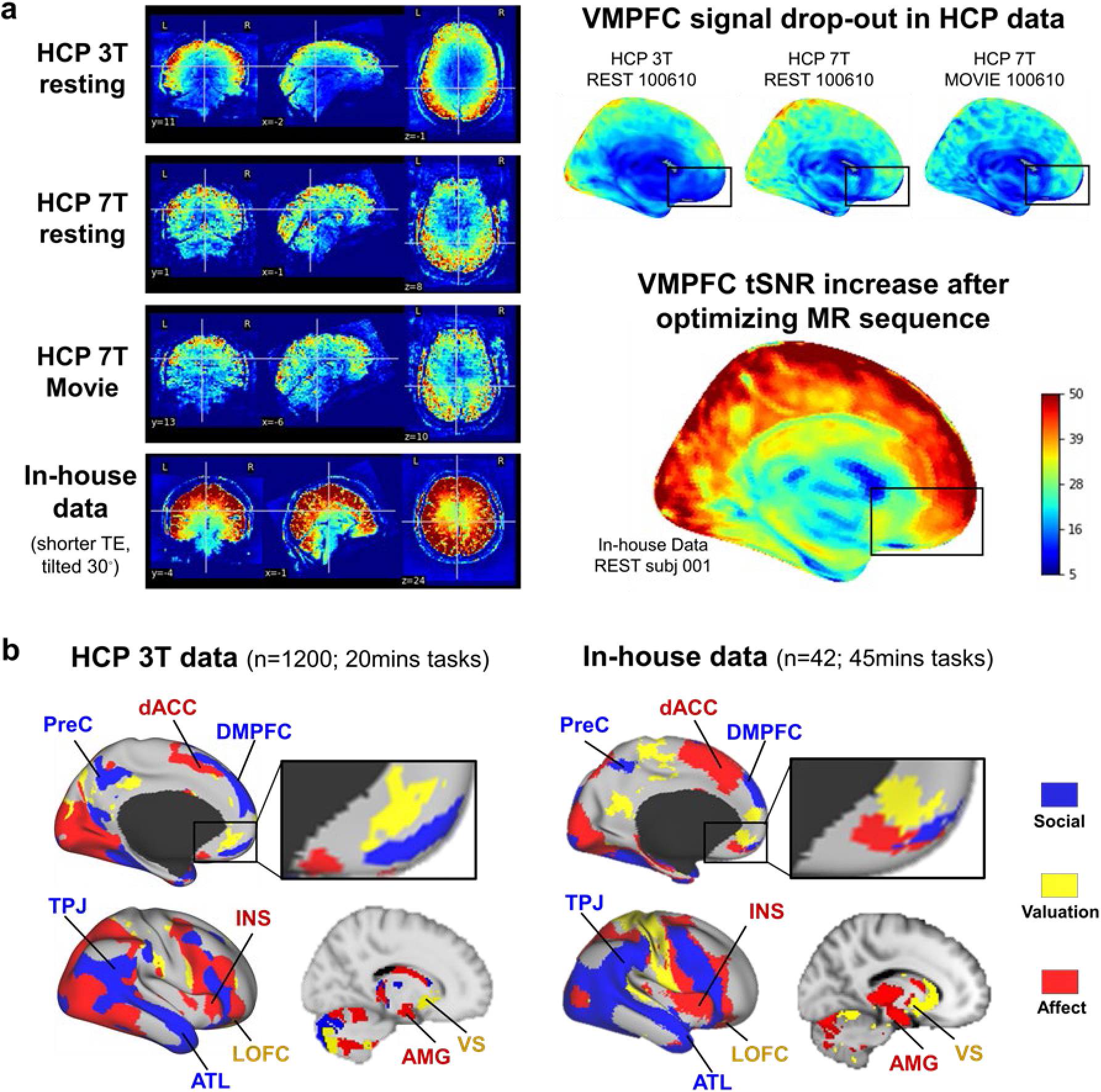
Sequence optimization restores VMPFC signal and reproduces the tripartite map in Study 3. **a**, Optimized acquisition (shorter TE and tilted 30 degrees) improves VMPFC’s data quality. Temporal SNR maps in our in-house dataset demonstrate substantial signal recovery in VMPFC (black bounding box in the right panel), which significantly outperforms HCP 3T and even HCP 7T (note that published reports of 7 T gains of ∼200-300% over 3T^102^ — underscoring that careful sequence optimization can offset susceptibility-induced signal loss in ventral frontal regions). **b,** Consistent functional topography with optimized data. Group-level winner-take-all maps from the in-house optimized dataset recapitulate the HCP pattern: posterior VMPFC aligns with the emotion circuitry (amygdala, insula, dACC), middle VMPFC with the reward system (VS, LOFC), and anterior VMPFC with the social brain network (PreC, TPJ, ATL, dmPFC). This confirms that the tripartite organization is not an artifact of sequence-specific signal loss and holds under improved ventral-frontal SNR. Notably, whereas the HCP Emotion task employed emotional face stimuli containing inherent social content, the in-house emotion task used emotionally salient IAPS scenes from which faces and social interactions were excluded, yet both paradigms converged on similar posterior VMPFC recruitment patterns. This convergence suggests that posterior VMPFC recruitment cannot be explained solely by the social nature of facial stimuli and instead reflects a broader affective processing tendency that generalizes across stimulus classes. All maps are shown in MNI space; color conventions match the main figures (blue = social, yellow = valuation, red = affect).

**Extended Data Fig. 5.**
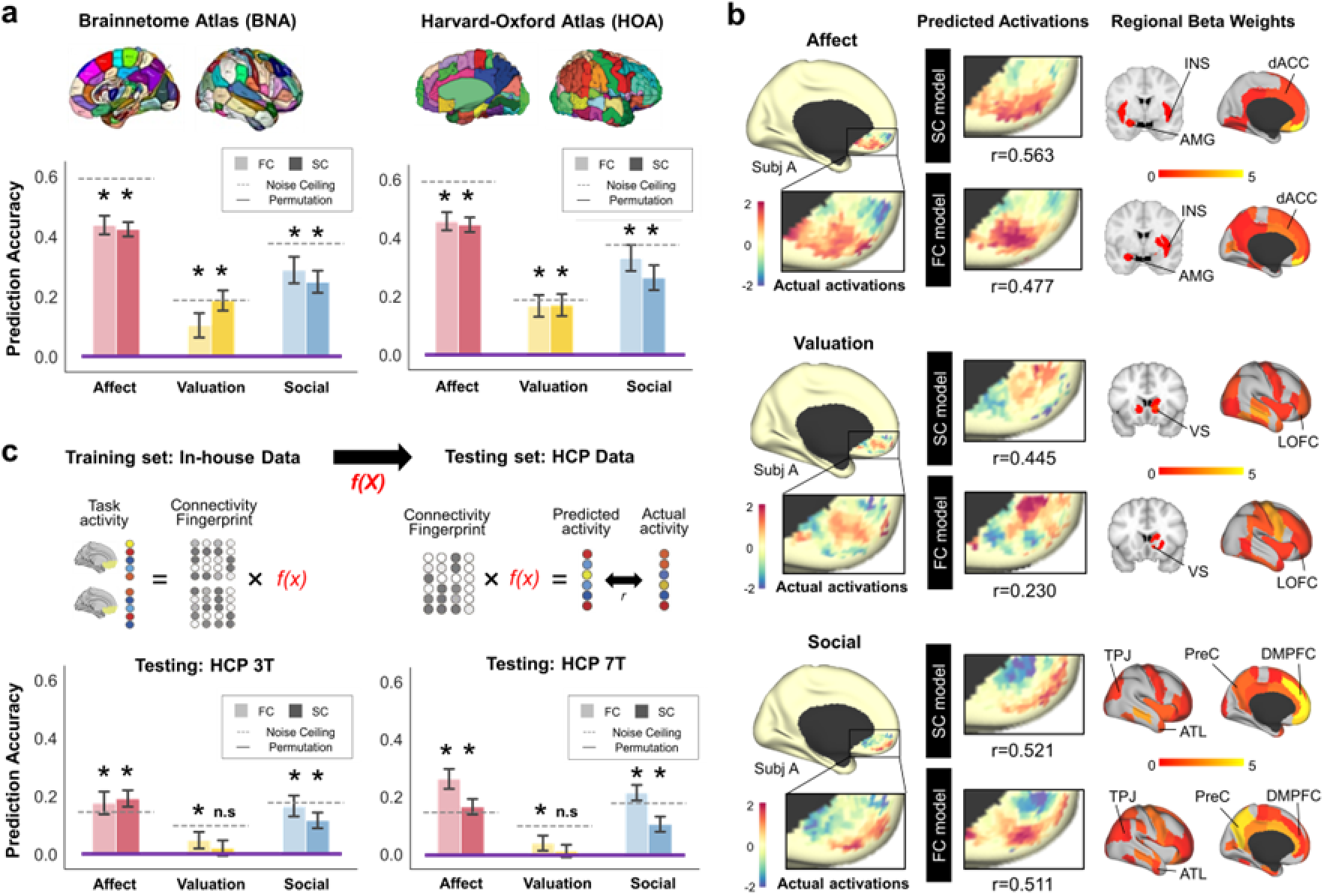
Generalization of connectivity-based prediction across brain atlases and datasets in Study 3. **a,** Main analysis of Study 3 (left) used whole-brain connectivity fingerprints defined by the BNA atlas (210 cortical + 36 subcortical parcels) together with the MDTB cerebellar parcellation (10 parcels). Repeating the same voxelwise connectivity-to-activation prediction using an alternative atlas set—Harvard-Oxford cortical atlas (96 parcels) combined with the Melbourne subcortical atlas (34 parcels) and MDTB cerebellar parcellation (10 parcels)—yields comparable prediction performance and similar connectivity projection motifs (see b), demonstrating that results are not dependent on atlas choice. **b,** Example predicted activation maps (SC, FC) and corresponding β-weight maps using the HOA atlas. Across atlases, connectivity features contributing most strongly to each VMPFC motif show a consistent pattern: posterior VMPFC is preferentially shaped by projections from the emotion circuitry (amygdala, insula, dACC); middle VMPFC by projections from the reward system (ventral striatum, lateral orbitofrontal cortex); and anterior VMPFC by projections from the social brain network (precuneus, temporoparietal junction, anterior temporal lobe, dorsomedial prefrontal cortex). **c,** Cross-dataset generalization. Predictive models trained on the in-house optimized dataset (with GLM weights fixed) were applied to independent HCP connectivity data (3T resting-state and 7T resting-state connectivity fingerprints) to predict task-evoked activation patterns within each VMPFC motif. Bars show out-of-sample prediction accuracy (Pearson r) for task contrasts associated with affect, valuation, and social domains, with noise ceilings (run 1 × run 2 reliability, dash line) and permutation baselines (purple line) shown for reference. Cross-dataset generalization is robust overall, with reduced performance for structural connectivity in the valuation condition. Color conventions for domains match main figures (red = affect, yellow = valuation, blue = social). Statistical procedures and model parameters are detailed in Methods.

**Extended Data Fig. 6.**
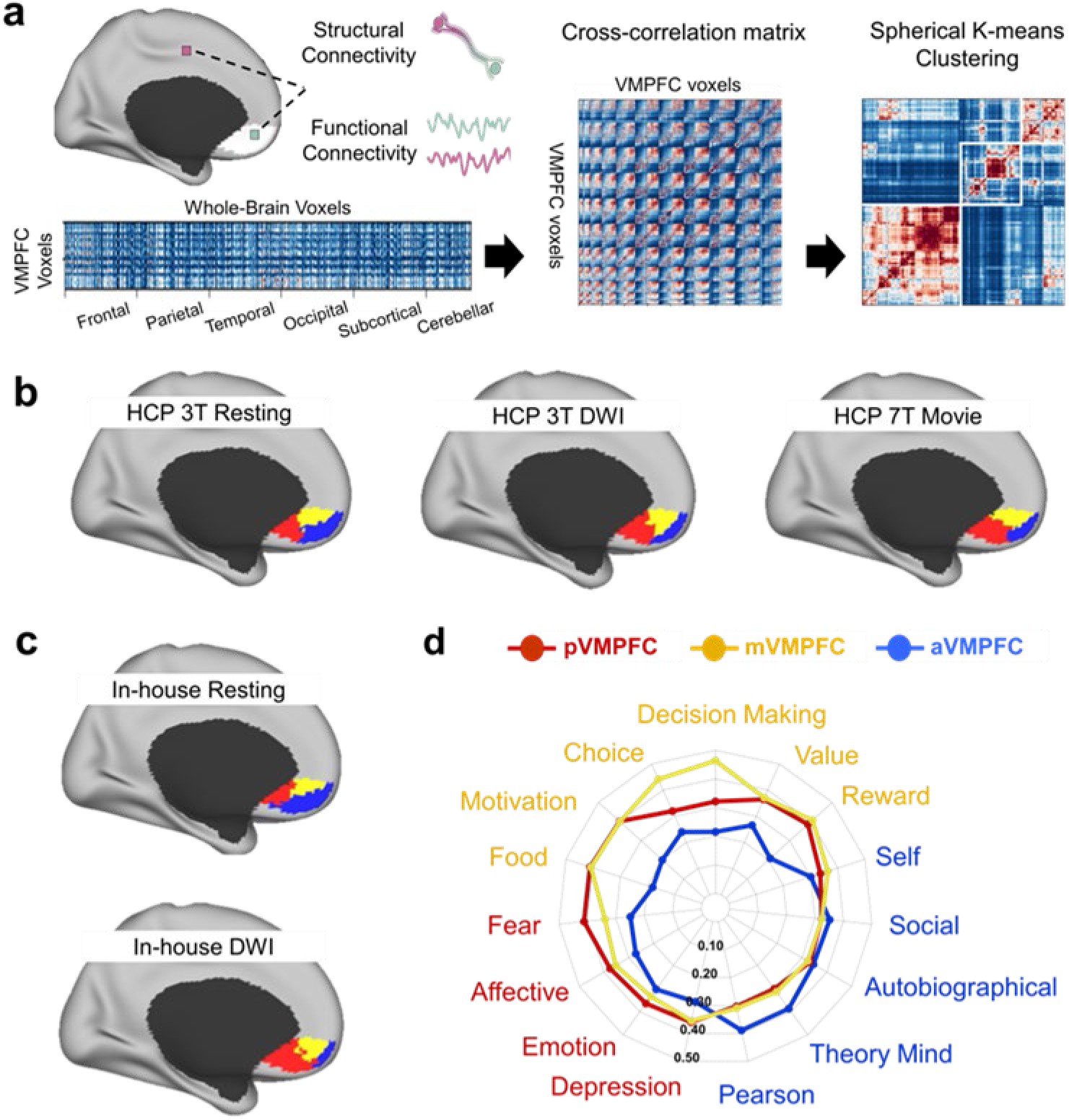
Connectivity-based parcellation of the VMPFC reveals a reproducible posterior–middle–anterior tripartition across datasets and modalities in Study 3. **a**, Parcellation pipeline. For each VMPFC voxel, we computed whole-brain functional connectivity (FC) from resting-state or movie fMRI and structural connectivity (SC) from diffusion tractography. Voxel-wise connectivity fingerprints were correlated to form a VMPFC similarity matrix, which was clustered using K-means to derive connectivity-defined functional motifs. **b,** HCP datasets. Connectivity-based parcellations derived from HCP-YA 3T resting FC and DWI SC, and HCP-YA 7T movie FC, each produced a consistent three-motif organization of the VMPFC, arranged along the anterior–posterior axis: aVMPFC (blue), mVMPFC (yellow), and pVMPFC (red). **c,** In-house datasets. The same posterior–middle–anterior tripartition was reproduced using in-house resting-state FC and DWI SC, confirming robustness across scanners, acquisition protocols, and participant cohorts. **d,** Functional decoding of connectivity-defined motifs. Term-based reverse-inference decoding of the three connectivity-defined VMPFC motifs, using the 15 a priori functional terms from Figure 1a, revealed motif-consistent functional specialization: the posterior motif with affect-related terms, the middle motif with valuation-related terms, and the anterior motif with social-related terms. Color conventions match all main figures (blue = social, yellow = valuation, red = affect). Parcellation and decoding procedures are described in Methods.

**Extended Data Fig. 7.**
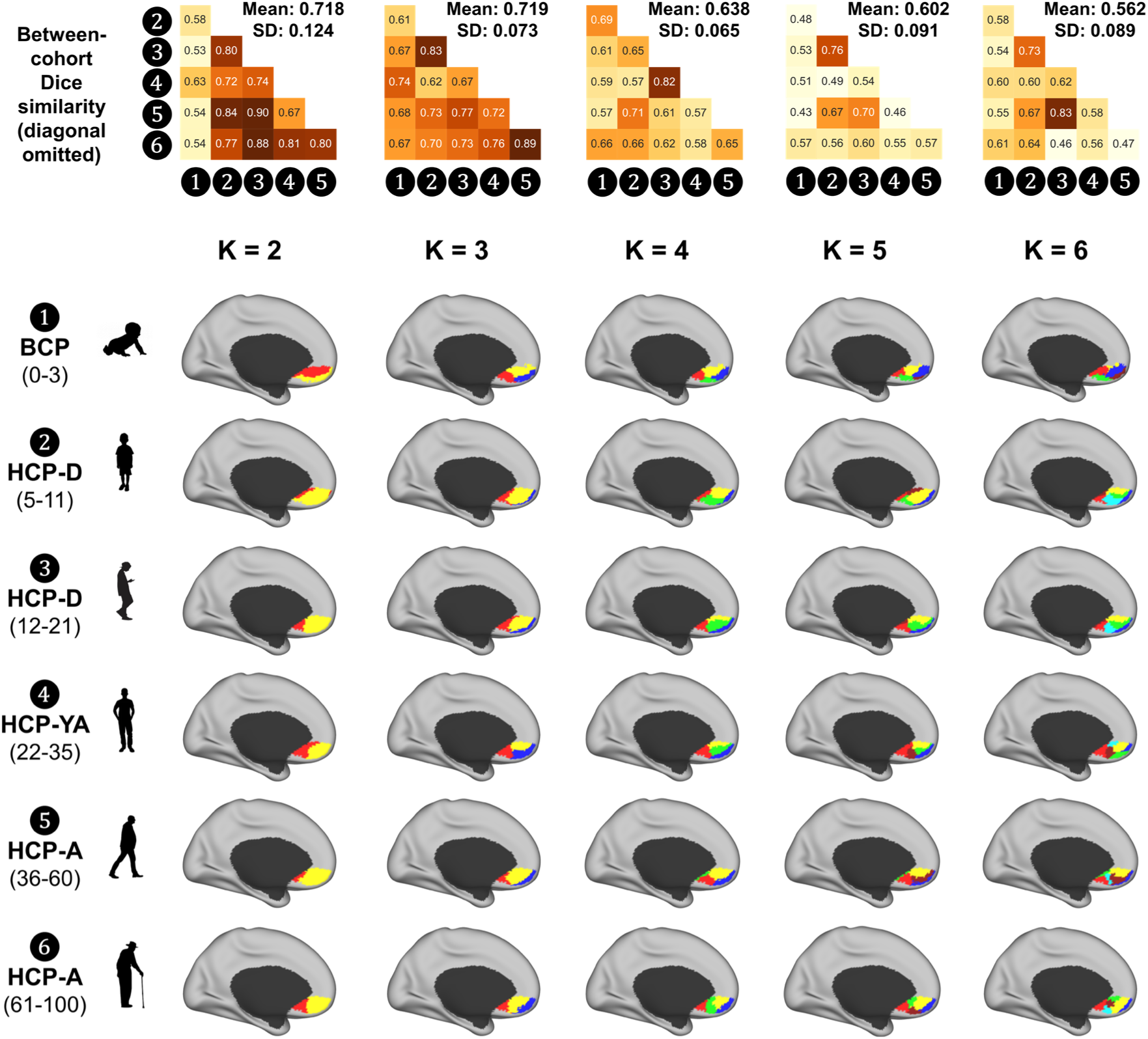
Cross-dataset agreement of VMPFC parcellations across cluster resolutions in Study 4. Top row, pairwise cross-cohort agreement matrices. For each cluster resolution (K = 2–6), heatmaps display the pairwise similarity between group-level VMPFC parcellations derived from different human lifespan cohorts (infant, child, adolescent, adult, and older adult). Rows and columns correspond to distinct cohorts; each cell reports the similarity between parcellations from two different cohorts, quantified using the macro-averaged Dice coefficient (macro-F1). Self-comparisons are excluded and therefore the diagonal is omitted. Numbers within cells indicate Dice values, and darker colors denote higher agreement. Mean and standard deviation (SD) of off-diagonal Dice values are shown above each matrix. Although K = 2 and K = 3 yield comparable mean similarity (0.718 vs. 0.719), K = 3 exhibits substantially lower variability across cohorts (SD = 0.073 vs. 0.124), indicating more uniform cross-dataset agreement. In contrast, K = 2 shows greater dispersion, driven primarily by reduced similarity involving the infant cohort (BCP). **Bottom rows, exemplar parcellations across cohorts and resolutions.** Cortical surface renderings show connectivity-defined VMPFC motifs for K = 2–6 in each cohort. At K = 3, all cohorts consistently exhibit a posterior–middle–anterior tripartition corresponding to the affect-, valuation-, and social-dominant motifs identified in Studies 1–3. In contrast, K = 2 yields less consistent spatial organization across cohorts, whereas K ≥ 4 produces finer subdivisions that fragment otherwise coherent motifs. Together, these results indicate that K = 3 provides the most stable and developmentally generalizable cross-dataset solution.

**Extended Data Fig. 8.**
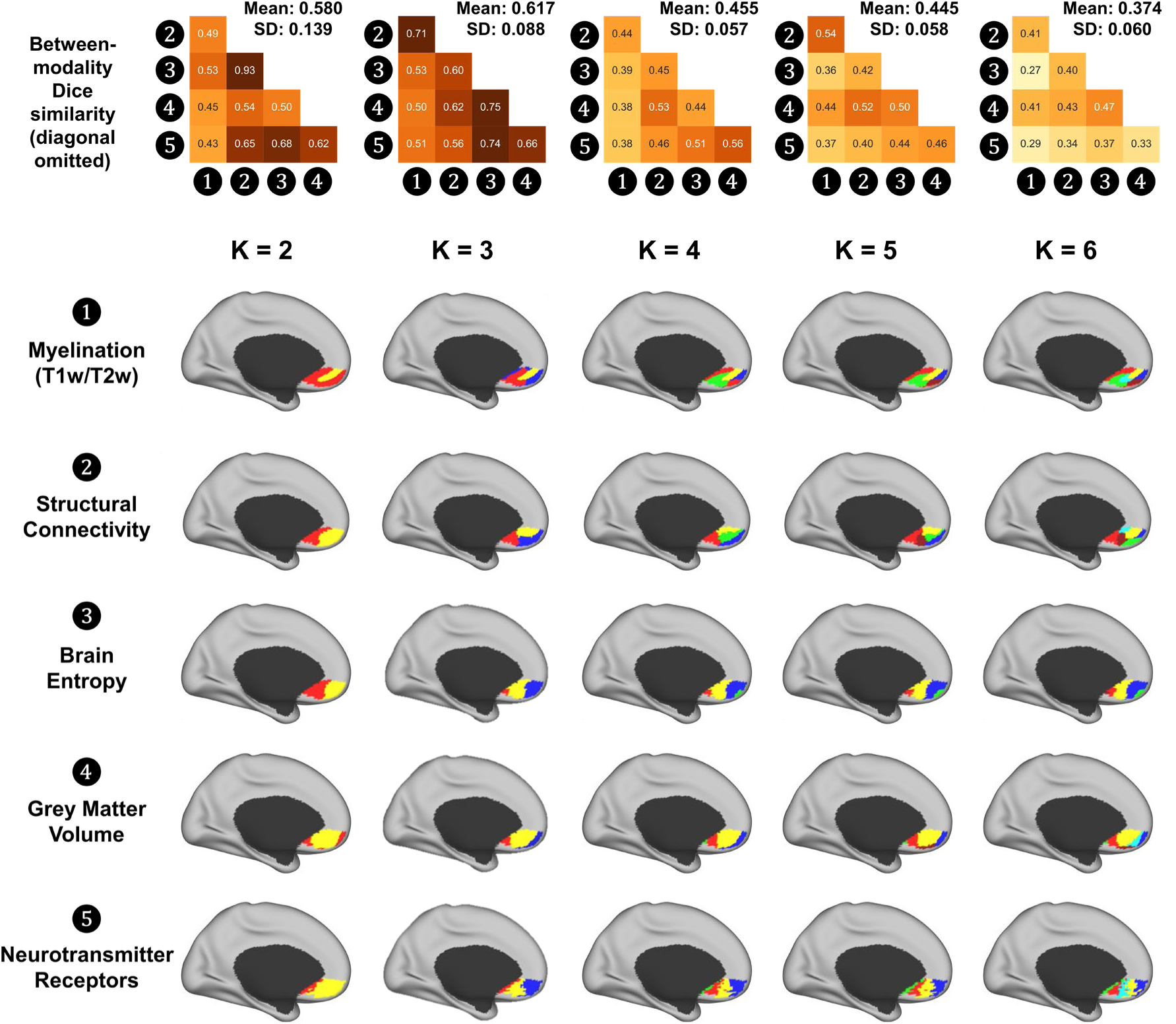
Cross-modality convergence identifies K=3 as the most stable VMPFC parcellation in Study 4. Top row, pairwise cross-modality agreement matrices. For each cluster resolution (K = 2-6), heatmaps display the pairwise similarity between VMPFC parcellations derived independently from different neurobiological modalities: cortical myelination (T1w/T2w) from HCP-YA structural MRI, structural connectivity (SC) from HCP-YA diffusion MRI, brain entropy (BEN) from HCP-YA resting-state fMRI, gray-matter volume (GMV) from HCP-YA structural MRI, and neurotransmitter receptor/transporter distributions (PET) from the Hansen Receptors Atlas. Rows and columns correspond to distinct modalities; each cell reports the similarity between parcellations obtained from two different modalities, quantified using the macro-averaged Dice coefficient (macro-F1). Self-comparisons (modality compared with itself) are excluded and therefore the diagonal is omitted. Mean and standard deviation (SD) of off-diagonal Dice values are shown above each matrix. Prior to computing Dice, parcel labels were optimally matched across modalities using one-to-one relabeling to account for arbitrary cluster indexing. Cell values indicate Dice coefficients, with darker colors denoting higher cross-modality agreement. **Bottom rows, modality-specific parcellations.** Group-level connectivity- or feature-defined VMPFC motifs derived independently from each modality are shown at the corresponding K. Colors follow the tripartite convention used throughout the manuscript (posterior/affect-dominant = red, middle/valuation-dominant = yellow, anterior/social-dominant = blue). Across modalities, K = 3 yields the highest mean off-diagonal Dice and is the only resolution for which all modality pairs achieve Dice ≥ 0.50, indicating that a three-motif solution provides the most consistent and biologically convergent partition of the human VMPFC. Other values of K show lower and more heterogeneous cross-modality agreement.

**Extended Data Fig. 9.**
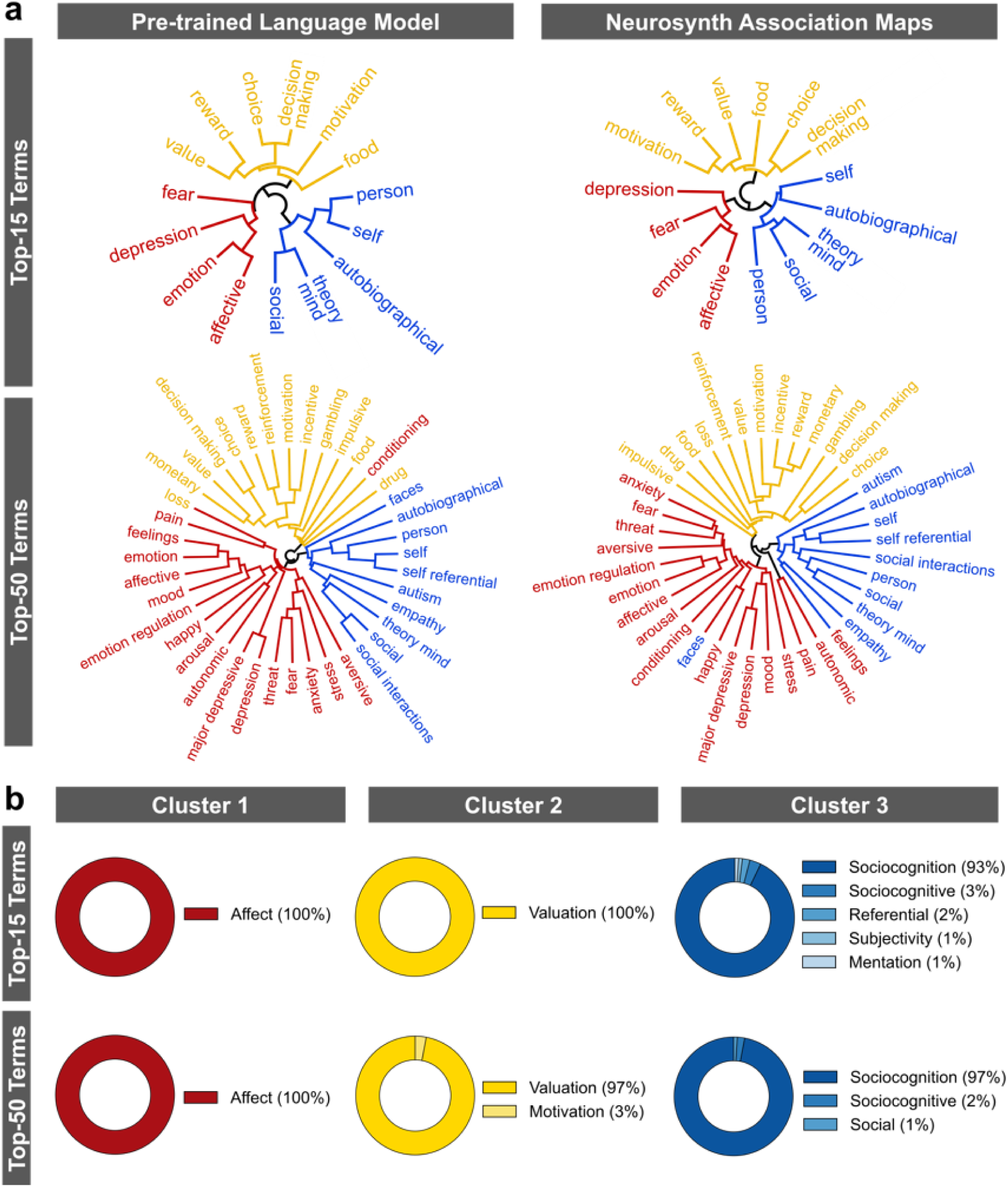
Data-driven identification and validation of the tripartite functional term structure in Study 1. **a**, Hierarchical clustering of cognitive terms in two independent representational spaces. The left column shows clustering based on semantic embeddings generated by a pretrained language model (Qwen3-Embedding-8B) using cosine distance. The right column shows clustering based on whole-brain Neurosynth association maps using correlation distance, with voxels inside the VMPFC mask excluded to avoid circularity with the VMPFC parcellation. Results are shown for both the top-15 terms used in the main analysis and an expanded set of 50 terms. Across both term sets and both representational spaces, terms consistently organize into three major branches corresponding to affective, valuation-related, and social domains. **b,** Independent automated labeling of the recovered clusters using a separate large language model (Gemini 3 Pro). Clusters were anonymized as “Cluster 1,” “Cluster 2,” and “Cluster 3,” and the model was provided only with the constituent terms. It was instructed to assign a single broad functional descriptor while avoiding anatomical network names or narrow task-specific labels. The procedure was repeated 100 times for both the 15-term and 50-term solutions. Donut plots show the proportion of runs assigning each label. Across both term sets, the dominant labels consistently corresponded to affect, valuation, and social cognition, providing convergent support for the tripartite functional interpretation of the VMPFC motifs.

**Extended Data Figure 10.**
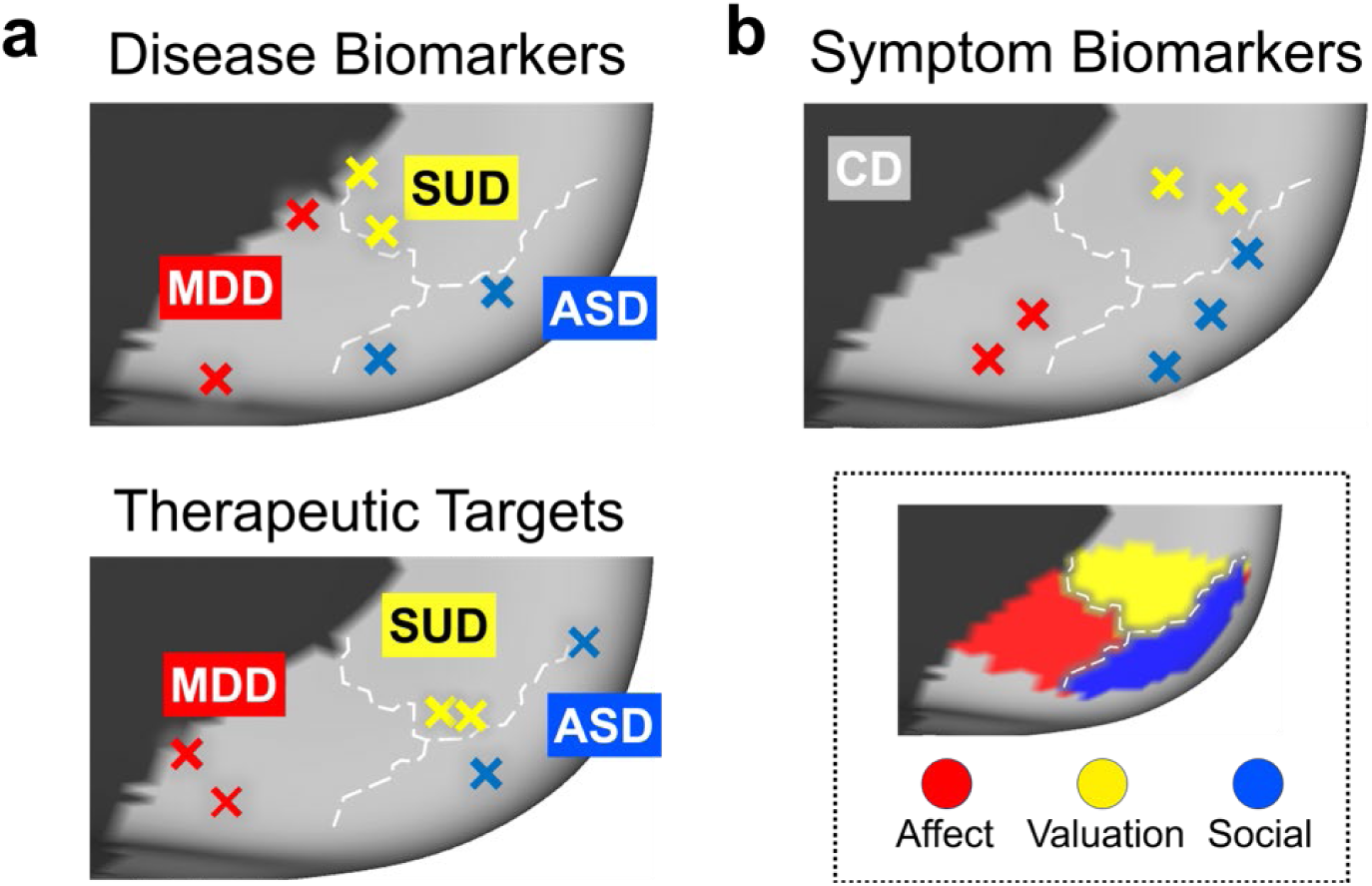
Exploratory clinical mapping of the VMPFC tripartite organization. a, Disorder-level mapping. Independent clinical findings associated with disease biomarkers (top) and therapeutic targets (bottom) were projected onto the VMPFC tripartite map. Illustrative disorders with predominantly affective (major depressive disorder, MDD; red), valuation-related (substance use disorder, SUD; yellow), or social (autism spectrum disorder, ASD; blue) dysfunction exhibit spatial distributions broadly consistent with the tripartite organization. MDD-related findings localize primarily to posterior VMPFC, SUD-related findings to middle VMPFC, and ASD-related findings to anterior VMPFC. **b, Symptom-level mapping within a heterogeneous disorder.** Peak coordinates associated with distinct symptom dimensions of conduct disorder (CD) were classified as affect-related (red), valuation-related (yellow), or social-related (blue). Symptom-related findings are distributed across all three VMPFC sectors, with different symptom dimensions showing preferential localization to posterior, middle, and anterior regions. The inset shows the reference tripartite organization identified in Study 1; dashed lines indicate motif boundaries. All coordinates are projected onto the medial surface of the left hemisphere for visualization. Detailed coordinates and source studies are provided in Supplementary Table 3.

## Supplementary Material

### Supplementary Methods

#### In-house Data with Optimized MRI Acquisition Details

All MRI data were collected at the Beijing Normal University Brain Imaging Center using a Siemens MAGNETOM Prisma 3T scanner equipped with a 64-channel head coil. Functional images were acquired using a T2-weighted gradient-echo echo-planar imaging (EPI)* sequence with multiband acceleration. To reduce susceptibility-induced signal dropout in ventral prefrontal regions, slices were acquired with a 30° tilt relative to the AC–PC line, providing improved signal coverage in the VMPFC. Each functional volume consisted of 72 axial slices (whole-brain coverage), with an in-plane resolution of 2 × 2 mm² and slice thickness of 2 mm (no gap). Acquisition parameters were: TR = 2000 ms, TE = 22 ms, flip angle = 90°, phase-encoding direction = anterior–posterior, FOV = 200 × 200 mm², bandwidth = 2380 Hz/Px, and multiband acceleration factor = 2.

A high-resolution T1-weighted structural image was acquired using a 3D MPRAGE sequence (voxel size = 1 × 1 × 1 mm³; TR = 2530 ms, TE = 2.27 ms, TI = 1100 ms, flip angle = 7°, FOV = 256 × 256 mm²). To correct for geometric distortions in the EPI data, a field map was acquired using a dual-echo spin-echo sequence (matched slice prescription; voxel size = 2 × 2 × 2 mm³; TR = 720 ms; TE₁ = 4.92 ms, TE₂ = 7.38 ms, flip angle = 60°). For diffusion MRI distortion correction, an additional reverse phase-encoded (PA) field map was acquired using the same parameters without diffusion weighting.

To evaluate the improvement afforded by this optimized acquisition protocol, temporal signal-to-noise ratio (tSNR) was computed using the Nipype confounds module (https://nipype.readthedocs.io). Compared to publicly available HCP data, the optimized protocol yielded substantially higher tSNR in the ventromedial prefrontal cortex (Extended Data Fig. 4), confirming the effectiveness of the acquisition adjustments for studying VMPFC function.

#### Non-Human Primate Dataset (PRIME-DE Macaque Oxford)

Non-human primate data were sourced from the Macaque Oxford dataset within the PRIMatE Data Exchange (PRIME-DE) initiative (Milham et al., 2018). Twenty rhesus macaques (Macaca mulatta) were scanned under light isoflurane anesthesia on a 3 T scanner using a four-channel coil; 19 animals passed preprocessing quality control (males, 4.01 ± 0.98 years, 6.61 ± 2.04 kg). T1w images were acquired with a 3D MPRAGE sequence (TR/TE = 2500/4.01 ms; FA = 8°; 0.5 mm isotropic resolution). Resting-state fMRI data were collected using a GRE-EPI sequence (TR/TE = 2000/19 ms; FA = 90°; 2 mm isotropic; 1600 volumes per animal; ∼53 min). Preprocessing followed Xu et al. (2018, 2019) including: structural denoising (Zuo & Xing, 2011), brain extraction with manual correction, tissue segmentation, and ANT-based normalization to the Yerkes19 template (Donahue et al., 2016). Functional data underwent despiking (AFNI 3dDespike), motion correction, global scaling, nuisance regression (WM, CSF, Friston-24 motion parameters), band-pass filtering (0.01–0.1 Hz), linear and quadric trend removal, coregistration to native space, volume-to-surface projection, and Gaussian smoothing (3 mm FWHM). Detailed protocols and QA information are available on the PRIME-DE portal (http://fcon_1000.projects.nitrc.org/indi/PRIME/oxford.html).

#### Multimodal Feature–Based Parcellation

To examine whether the tripartite organization of the ventromedial prefrontal cortex (VMPFC) could be recovered from imaging modalities beyond task activity and brain connectivity, we employed four independent profile-based analyses: gray-matter volume (GMV), cortical myelination, resting-state brain entropy (BEN), and neurotransmitter receptor/transporter distribution. Each modality was processed independently within its respective cohort, and voxel- or vertex-wise feature covariance profiles were used for clustering analyses (K = 2–6, see Extended Data Fig. 8).

##### Gray-Matter Volume (GMV)

A profile-based approach using inter-individual variations in GMV was applied separately to HCP-YA dataset. Modulated GMV maps were derived from participants’ T1-weighted images following the standard voxel-based morphometry (VBM) framework (Ashburner & Friston, 2000). This included tissue segmentation, spatial normalization to the MNI152NLin2009cAsym template, and modulation using Jacobian determinants to preserve local volumetric information. We additionally followed the FSLVBM pipeline, creating a study-specific GM template (fslvbm_2_template) and non-linearly registering individual GM maps using fslvbm_3_proc. Spatial smoothing employed a Gaussian kernel (FWHM = 6 mm for HCP-YA). We computed each VMPFC voxel’s covariance with every brain voxel across participants, then calculated a voxel-by-voxel cross-correlation matrix describing the similarity of whole-brain GMV covariance profiles within the VMPFC. The resulting matrices were clustered using k-means (K = 2–6); cluster validity was assessed via Silhouette and Davies–Bouldin indices. Guided by prior hypotheses and meta-analytic evidence, K = 3 was selected as the optimal solution.

##### Cortical Myelination

A second approach used cortical myelination (T1w/T2w ratio) to define VMPFC subregions in the HCP-YA dataset. Myelin maps were obtained from the standard release file MyelinMap_BC.32k_fs_LR.dscalar.nii distributed with the HCP-YA dataset. Using Connectome Workbench (v2.0.1), the volumetric VMPFC mask (defined in MNI space) was projected to the 32k_fs_LR surface and separated into left and right hemispheres. For each participant, vertex-wise myelination index values within the VMPFC were extracted to form subject-specific feature vectors. Covariance profiles were then computed across participants for each vertex relative to the entire cortical surface, yielding a group-level vertex-by-vertex similarity matrix. This matrix was clustered using k-means (K = 2–6), yielding surface-based myelin-derived subregions.

##### Brain Entropy (BEN)

Another profile-based analysis leveraged BEN to delineate VMPFC subregions. BEN quantifies the irregularity and complexity of spontaneous brain activity (Wang et al., 2014). Analyses were conducted on HCP-YA dataset. For each participant, BEN maps were generated from preprocessed resting-state fMRI data using the BENtbx toolbox with a window length = 3 and a cutoff threshold = 0.6 (Wang et al., 2014). HCP-YA fMRI data followed the consortium’s minimal preprocessing, then underwent additional band-pass filtering (0.01–0.1 Hz) and Gaussian smoothing (FWHM = 6 mm). BEN maps were smoothed with a 6 mm Gaussian kernel. We computed voxel-wise covariance between each VMPFC voxel’s BEN profile and all brain voxels, constructed a cross-correlation matrix of these covariance vectors, and clustered it with k-means (K = 2–6).

##### Neurotransmitter Receptor and Transporter Profiles

A fourth analysis used spatial distributions of neurotransmitter receptors and transporters, based on the rationale that subregions with similar neurochemical signatures may share functional roles. We obtained receptor/transporter density maps from the Hansen Receptors Atlas (i.e., NetNeurolab open repository, https://github.com/netneurolab/hansen_receptors/tree/main/data/PET_nifti_image), comprising 19 PET-derived maps spanning nine major neurotransmitter systems aggregated from > 1,200 healthy individuals (Hansen et al., 2022). Maps were resampled to 3 mm isotropic resolution; one failed resampling, leaving 18 usable maps. To focus on cortical neurochemistry and reduce subcortical noise, values were restricted to the cortical sheet and parcellated using the 1,000-region Schaefer atlas (Schaefer et al., 2018) implemented in the Neuromaps toolbox (Markello et al., 2022). For each cortical parcel, an 18-dimensional receptor/transporter profile vector was computed by averaging density values across the 18 maps. These parcel-wise profiles were used to estimate cross-parcel similarity within the VMPFC, and the resulting correlation matrices were clustered using k-means (K = 2-6).

**Supplementary Table 1.**
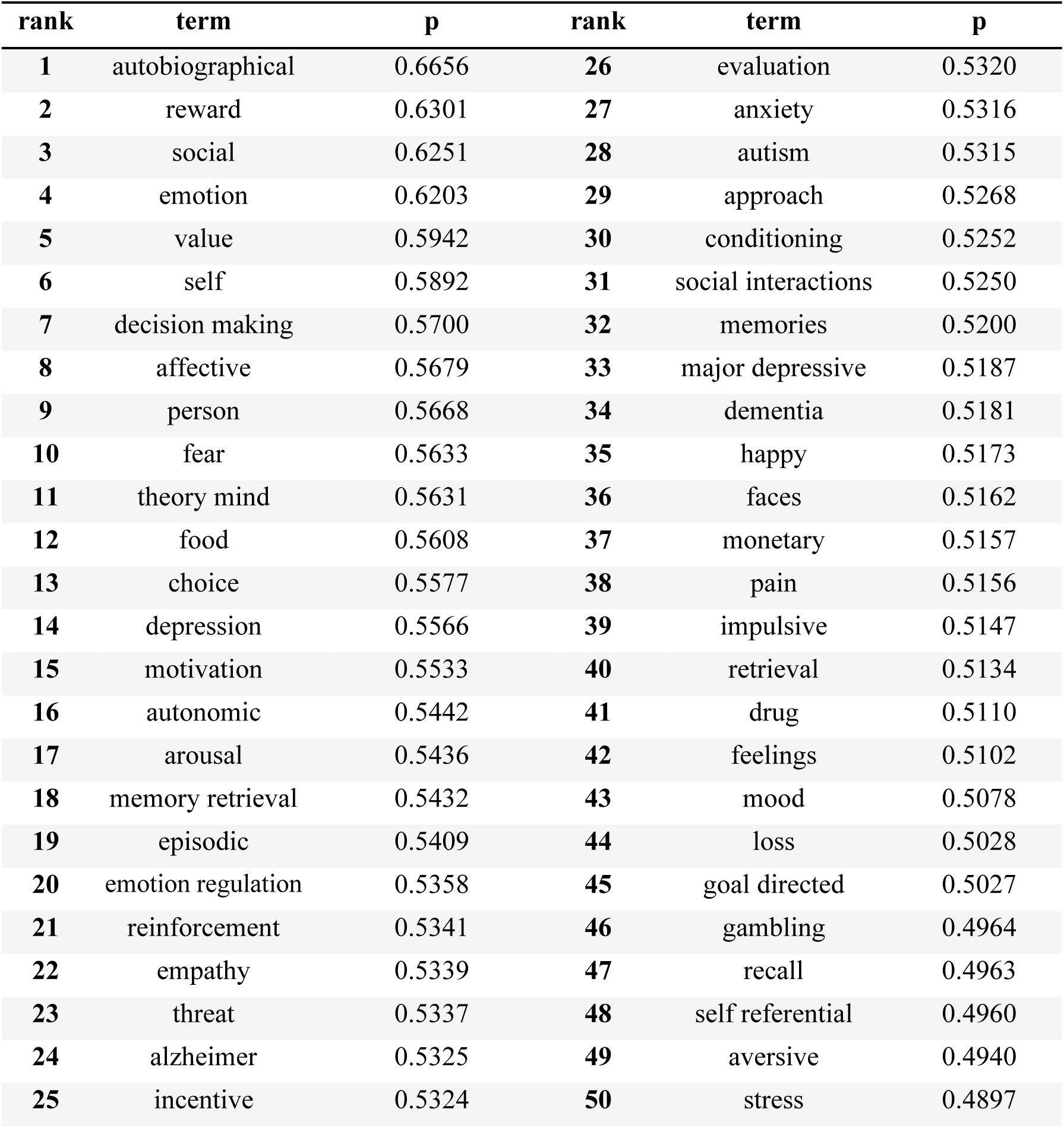
Meta-analytic reverse inference decoding of the VMPFC using NS+ (top 50 terms) in Study 1. The column p denotes as reverse-inference posterior probability (not a significance threshold) under a uniform prior probability = 0.5 used for ranking. Top 15 terms were showed in Fig. 1a.

**Supplementary Table 2.** Image Databases Used for ANN simulation and validation in Study 2.

| Domains | Datasets | No. Images | Key Contents | URL and References |
| --- | --- | --- | --- | --- |
| **Social** | WIDER-Parade | 1,150 | Social parade scenes<br>(unable to see people's facial expression) | http://shuoyang1213.me/WIDERFACE<br>Yang, S. et al. (2016). WIDER FACE: A face detection benchmark. CVPR, 5525–5533. |
|  | PISC<br>(People in Social Context) | 22,690 | People in Social Context<br>(unable to see people's facial expression) | https://zenodo.org/record/1059155<br>Li, W. et al. (2017). People in Social Context: Recognizing people's roles in social relations. CVPR, 1354–1362. |
| **Valuation** | Food-5K | 2,500 | Diverse Food items | https://mmspg.epfl.ch/downloads/food-image-datasets/<br>Singla, A. et al. (2016). Food/non-food image classification and food categorization using pre-trained GoogLeNet. MADiMa, 3–11. |
|  | Food-11 | 3,430 | Diverse Food items | https://mmspg.epfl.ch/downloads/food-image-datasets/<br>Singla, A. et al. (2016). Food/non-food image classification and food categorization using pre-trained GoogLeNet. MADiMa, 3–11. |
| **Affect** | EXCEED<br>(Extreme Climate Event Dataset) | 100 | Extreme climate event images<br>(floods, droughts) | https://doi.org/10.6084/m9.figshare.4602016.v1<br>Magalhães, R. et al. (2018). The EXCEED database: Extreme climate event dataset for affective computing. Figshare. |
|  | GAPED | 415 | Negative affective and emotional scenes | https://www.unige.ch/cisa/research/materials-and-online-research/research-material/<br>Dan-Glauser, E. & Scherer, K. R. (2011). The Geneva Affective Picture Database (GAPED): A new resource for research on emotion and attention. Behavior Research Methods, 43(2), 468–477. |
| **Social**<br>**×**<br>**Value** | CFD<br>(Chicago<br>Face<br>Database) | 108 | Standardized<br>human faces<br>with<br>attractiveness<br>ratings<br>(only faces<br>with neutral<br>emotions<br>selected) | https://chicagofaces.org<br>Ma, D. S. et al. (2015). The Chicago Face<br>Database: A free stimulus set of faces and<br>norming data. Behavior Research<br>Methods, 47(4), 1122–1135. |
| **Value**<br>**×**<br>**Affect** | Antique<br>and aesthetic<br>object<br>dataset | 100 | Images of<br>antique<br>artworks<br>evoking<br>aesthetic<br>emotion | Self-curated collection from open-access<br>museum image repositories. |
| **Social**<br>**×**<br>**Affect** | SMID | 731 | Socio-Moral<br>Image<br>Database | https://osf.io/jd6uc/<br>Crone, D. L. et al. (2018). The Socio-Moral Image Database (SMID): A novel<br>stimulus set for moral psychology. PLoS<br>ONE, 13(1), e0190954. |
| **Social**<br>**×**<br>**Value**<br>**×**<br>**Affect** | NAPS_ERO<br>(Nencki<br>Affective<br>Picture<br>System) | 200 | Erotic subset<br>of NAPS | https://naps.nencki.gov.pl/<br>Marchewka, A. et al. (2014). The Nencki<br>Affective Picture System (NAPS):<br>Cross-cultural validation and content<br>description. Behavior Research Methods,<br>46(2), 596–610. |

**Supplementary Table 3.**
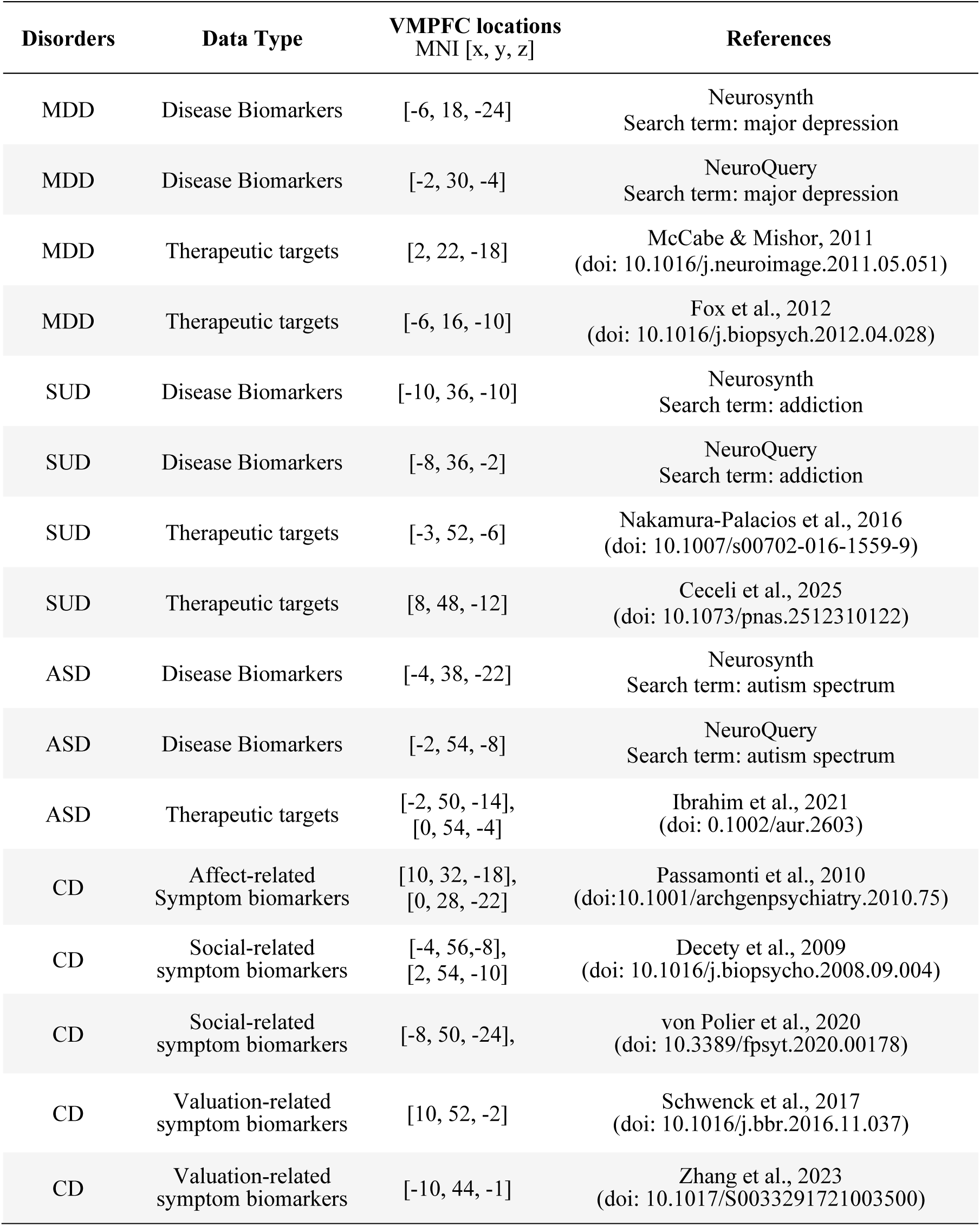
Coordinates and source studies used for exploratory clinical mapping of the VMPFC tripartite organization.

### Supplementary Figures

**Supplementary Figure 1.**
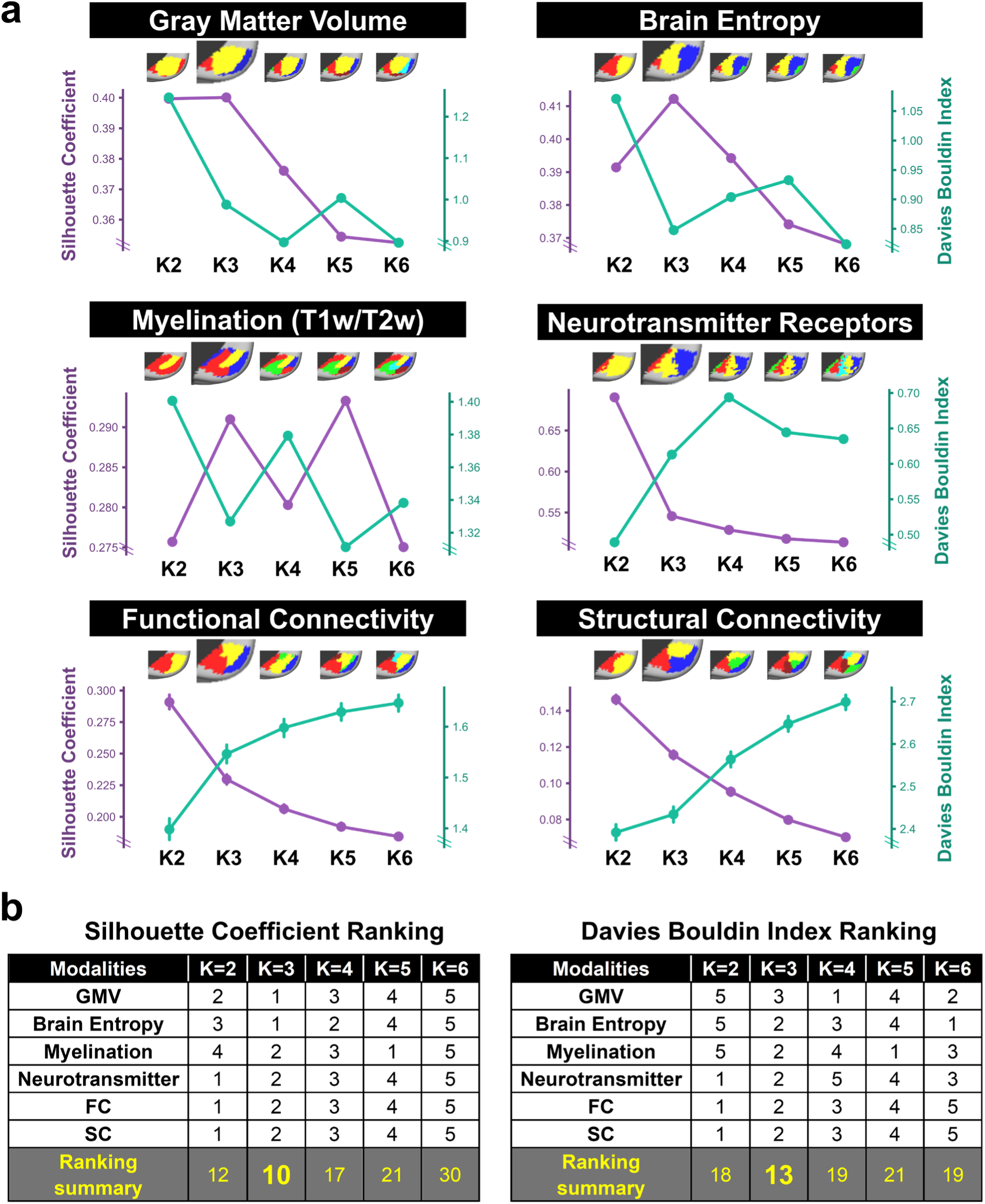
Internal clustering metrics across neurobiological modalities and consensus selection of K = 3 in Study 4. a, Modality-specific clustering solutions. Silhouette coefficients (purple; higher values indicate better cluster separation) and Davies–Bouldin indices (green; lower values indicate better clustering) are shown for K = 2–6 across six independent modalities: gray-matter volume covariance, brain entropy, myelination (T1w/T2w), neurotransmitter receptor density, functional connectivity, and structural connectivity. Representative VMPFC parcellations are displayed above each metric profile. Different modalities favor different resolutions, indicating sensitivity to distinct spatial scales of cortical organization. Connectivity-based measures preferentially support coarser partitions (K = 2), consistent with their emphasis on large-scale network architecture, whereas microstructural measures such as myelination favor finer subdivisions (K = 5), reflecting greater local differentiation in cortical properties. Gray-matter volume and brain entropy show optima near K = 3, consistent with an intermediate organizational scale. Such variation is expected because different neurobiological features capture partially overlapping but non-identical aspects of cortical architecture. **b, Cross-modal rank aggregation.** For each modality, K = 2–6 were ranked according to the Silhouette coefficient and Davies–Bouldin index separately, and ranks were summed across modalities. Lower values indicate better overall performance. K = 3 achieved the best aggregate ranking under both metrics, demonstrating that it is the most consistently supported solution across modalities. Combined with its superior cross-dataset reproducibility (Extended Data Fig. 7), maximal cross-modality convergence (Extended Data Fig. 8), and correspondence to the affective, valuation, and social motifs identified in Studies 1–3, these results support K = 3 as the strongest cross-modal consensus description of VMPFC organization.

**Supplementary Figure 2.**
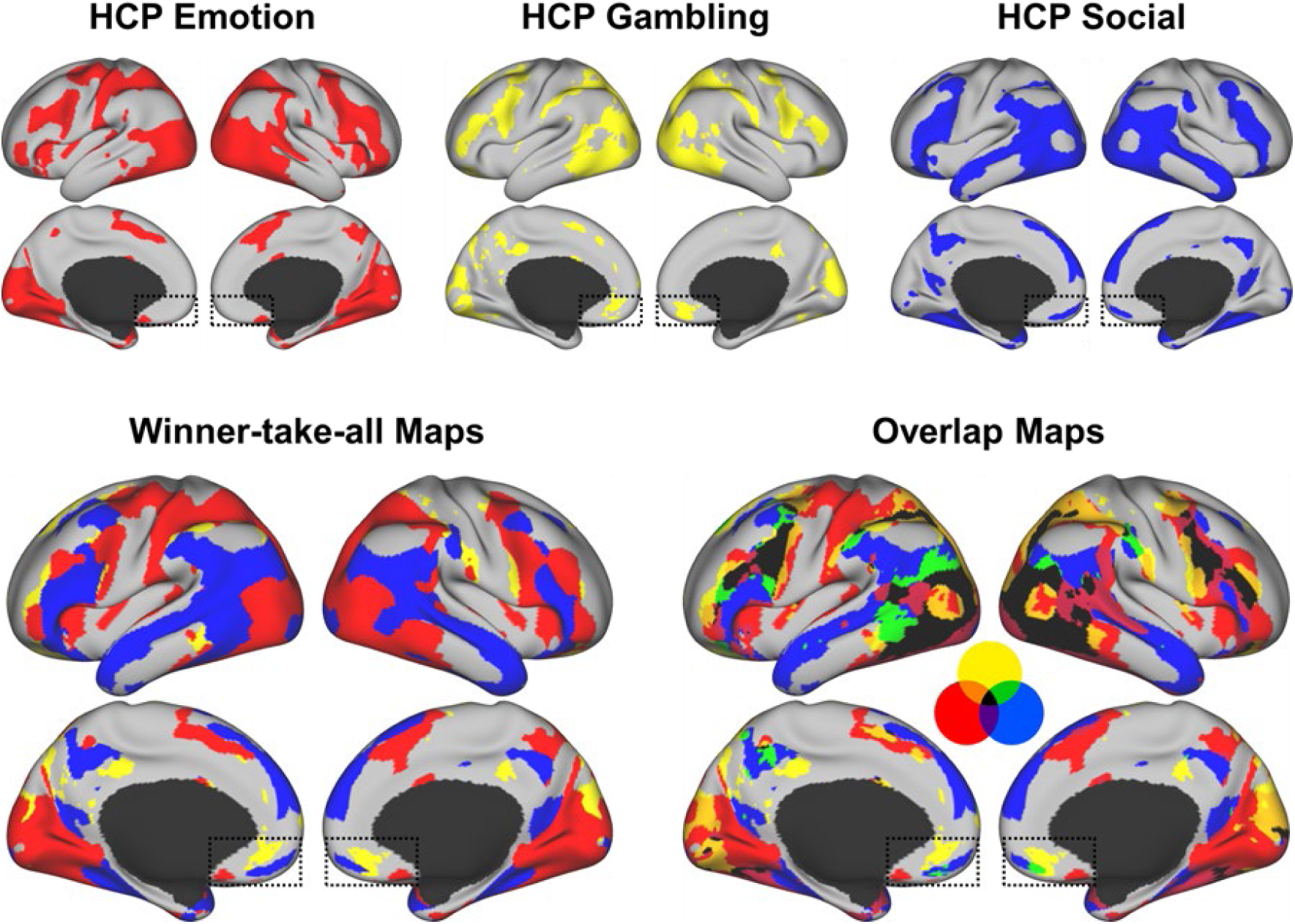
Standard activation, winner-take-all, and overlap maps for HCP task domains in Study 2. **Top row** shows independent thresholded group-level activation maps for the Affect (Emotion), Valuation (Gambling), and Social task contrasts used in Fig. 2. Maps are thresholded at Cohen’s d > 0.2 and displayed separately for each domain. **Bottom left** shows the winner-take-all (WTA) map, in which each voxel is assigned to the domain with the largest Cohen’s d value. This visualization highlights dominant functional preferences and reveals a posterior–middle–anterior ordering within the VMPFC (dashed boxes). **Bottom right** shows the overlap map derived from the thresholded activation maps. Colors indicate voxels activated by one or more domains, demonstrating substantial overlap among affective, valuation-related, and social activations, particularly within association cortex. Despite this overlap, the posterior–middle–anterior ordering remains evident, indicating that the tripartite organization is not an artifact of the WTA procedure. Together, these results suggest that the VMPFC is organized into partially overlapping functional motifs embedded within an integrative architecture rather than into rigidly segregated modules

**Supplementary Figure 3.**
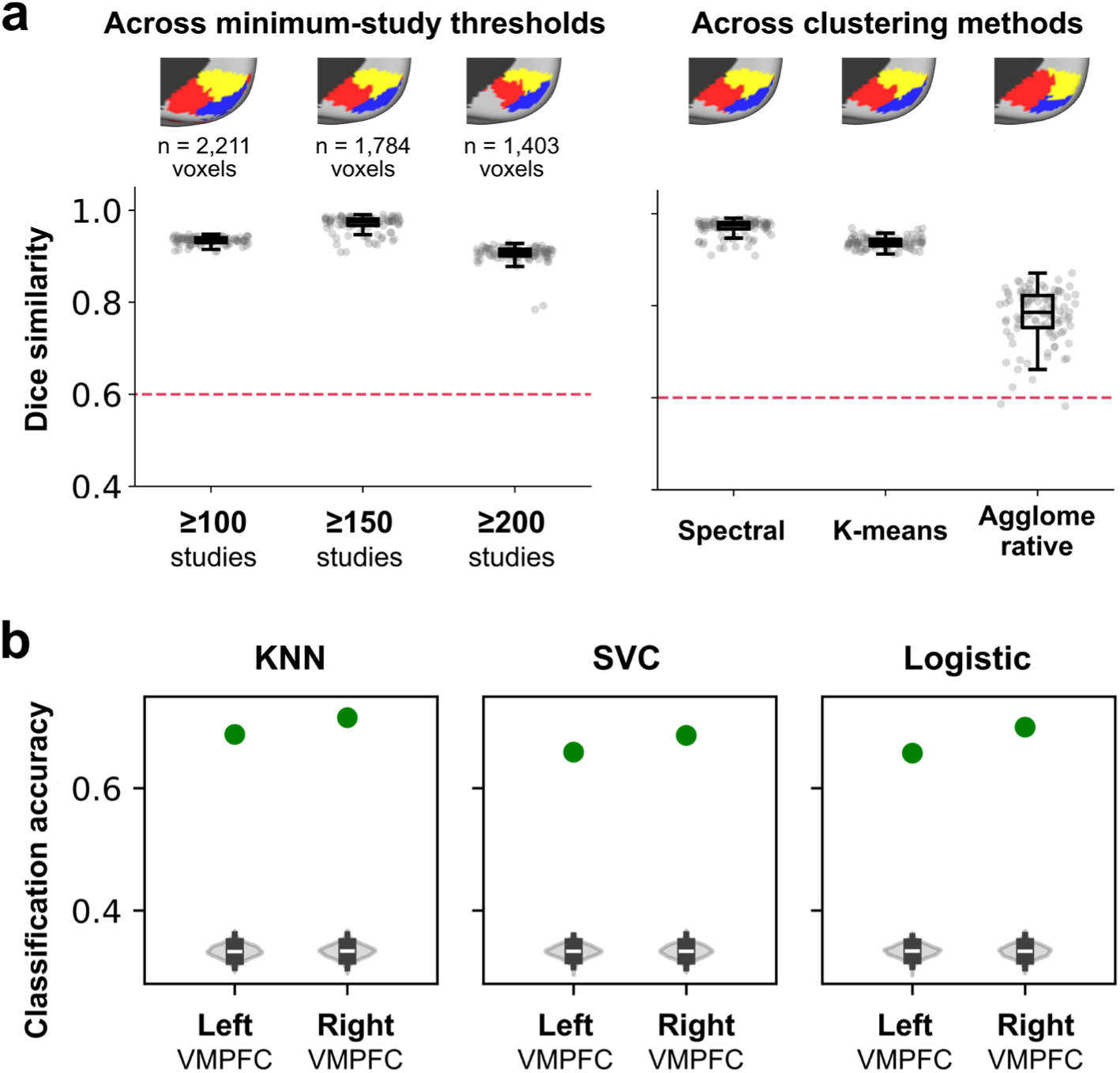
Robustness of the tripartite VMPFC organization across clustering parameters, algorithms, and classifiers in Study 1 and Study 2. **a,** Stability of the K = 3 parcellation across voxel-selection criteria and clustering algorithms. *Left,* spatial similarity (Dice coefficient) of the tripartite VMPFC solution across 100 independent K-means runs under different minimum-study inclusion thresholds used for MACM (≥100, ≥150, and ≥200 studies per voxel). *Right,* comparison of parcellation stability across three clustering algorithms: Spectral Clustering, K-means, and Agglomerative Clustering. Grey dots represent individual runs; boxplots show medians and interquartile ranges. The representative brain maps displayed above each condition correspond to the solution exhibiting the lowest spatial similarity (smallest Dice coefficient) among the 100 repetitions, providing a conservative estimate of robustness. The red dashed line indicates a reference similarity threshold (Dice = 0.60). Across all thresholds and algorithms, the tripartite posterior–middle–anterior organization was consistently recovered, indicating that the reported parcellation reflects a stable feature of the underlying co-activation structure rather than an artifact of a specific clustering procedure. **b,** Robustness of individual-level classification across classifier families. The subject-level motif classification analysis was repeated using three supervised learning algorithms: K-nearest neighbors (KNN), Support Vector Classification (SVC), and Logistic Regression. Green dots indicate observed classification accuracy for the left and right VMPFC. Grey violin plots represent null distributions obtained from permutation testing. All classifiers achieved accuracies substantially above chance in both hemispheres, demonstrating that individual-level validation of the tripartite organization is robust to classifier choice.

**Supplementary Figure 4.**
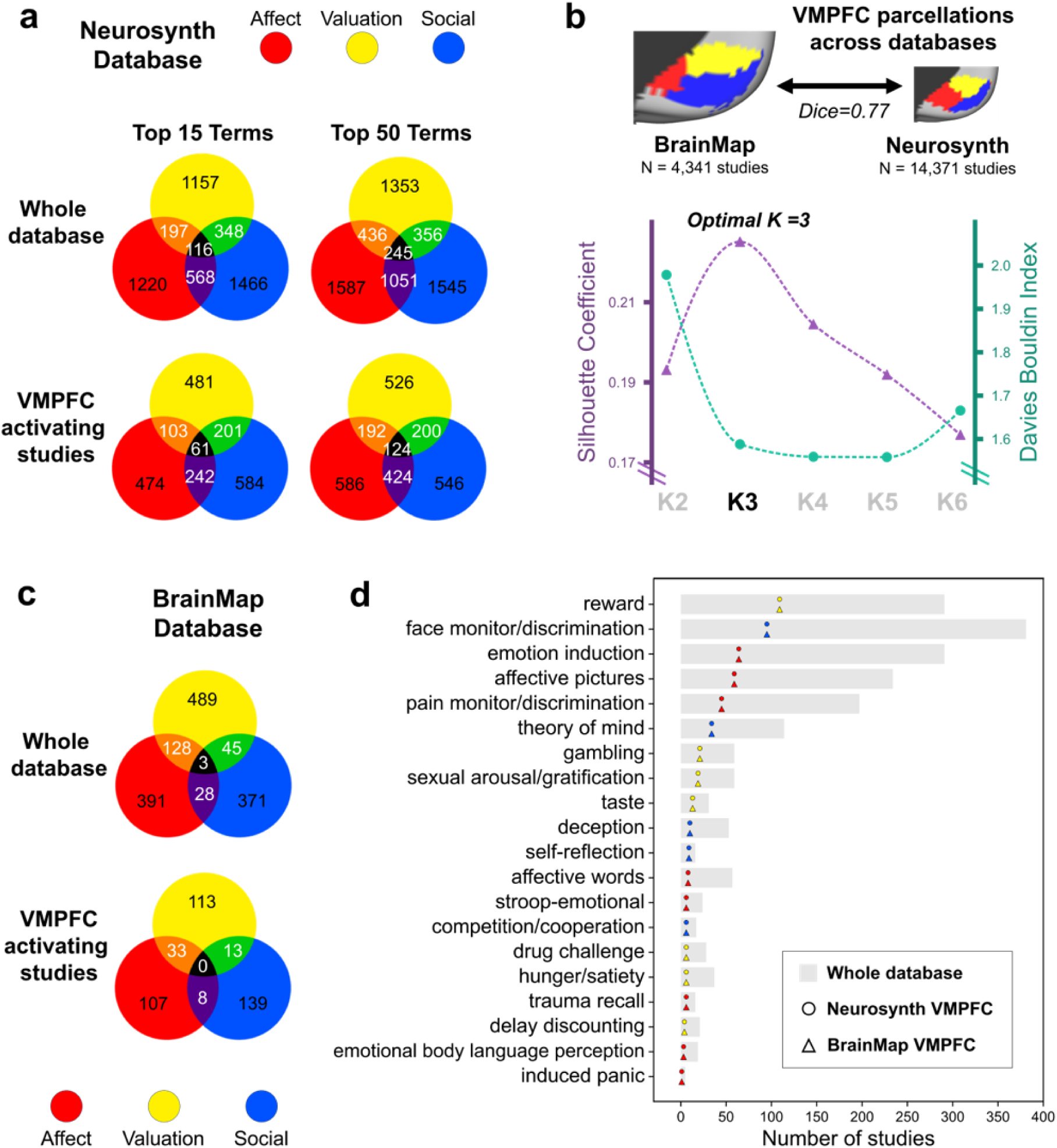
Distribution of studies across functional domains and experimental paradigms, and cross-database validation of the VMPFC parcellation in Study 1. **a,** Distribution of studies across functional domains in Neurosynth. Counts are shown for studies associated with the top 15 (left) and top 50 (right) VMPFC-related terms. The top row shows counts across the entire Neurosynth database, whereas the bottom row shows counts restricted to studies contributing to the VMPFC meta-analysis. Across both term sets and both study subsets, affective-, valuation-, and social-related studies are represented at broadly comparable frequencies, with substantial overlap between domains. **b,** Cross-database validation of the VMPFC parcellation. Independent MACM analyses performed in BrainMap (N = 4,341 studies) and Neurosynth (N = 14,371 studies) recovered highly similar tripartite VMPFC organizations (Dice coefficient = 0.77). Cluster-validity indices such as the Silhouette coefficient (purple) peaks at K = 3, while the Davies–Bouldin index (green) is comparably low across K = 3–5, supporting a three-cluster solution. Insets show the BrainMap- and Neurosynth-derived K = 3 parcellations. **c,** Distribution of studies across functional domains in BrainMap. Counts are shown for all BrainMap studies (top) and for studies contributing to the BrainMap-derived VMPFC parcellation (bottom). Similar to Neurosynth, the three domains occur at broadly comparable frequencies, with no single domain accounting for a majority of VMPFC-activating studies. **d,** Distribution of experimental paradigms in BrainMap. Gray bars indicate the total number of studies associated with each paradigm across the entire BrainMap database. Colored circles indicate the number of studies activating the Neurosynth-derived VMPFC, and colored triangles indicate the number of studies activating the BrainMap-derived VMPFC. Paradigms are color-coded according to their dominant functional domain (red = Affect, yellow = Valuation, blue = Social). No single paradigm dominates the VMPFC literature; the most frequent paradigm (i.e., “reward”) accounts for only 13.20% of studies in the Neurosynth VMPFC and 13.49% of studies in the BrainMap VMPFC, indicating that the observed tripartite organization is unlikely to be driven by overrepresentation of any individual task type.

**Supplementary Figure 5.**
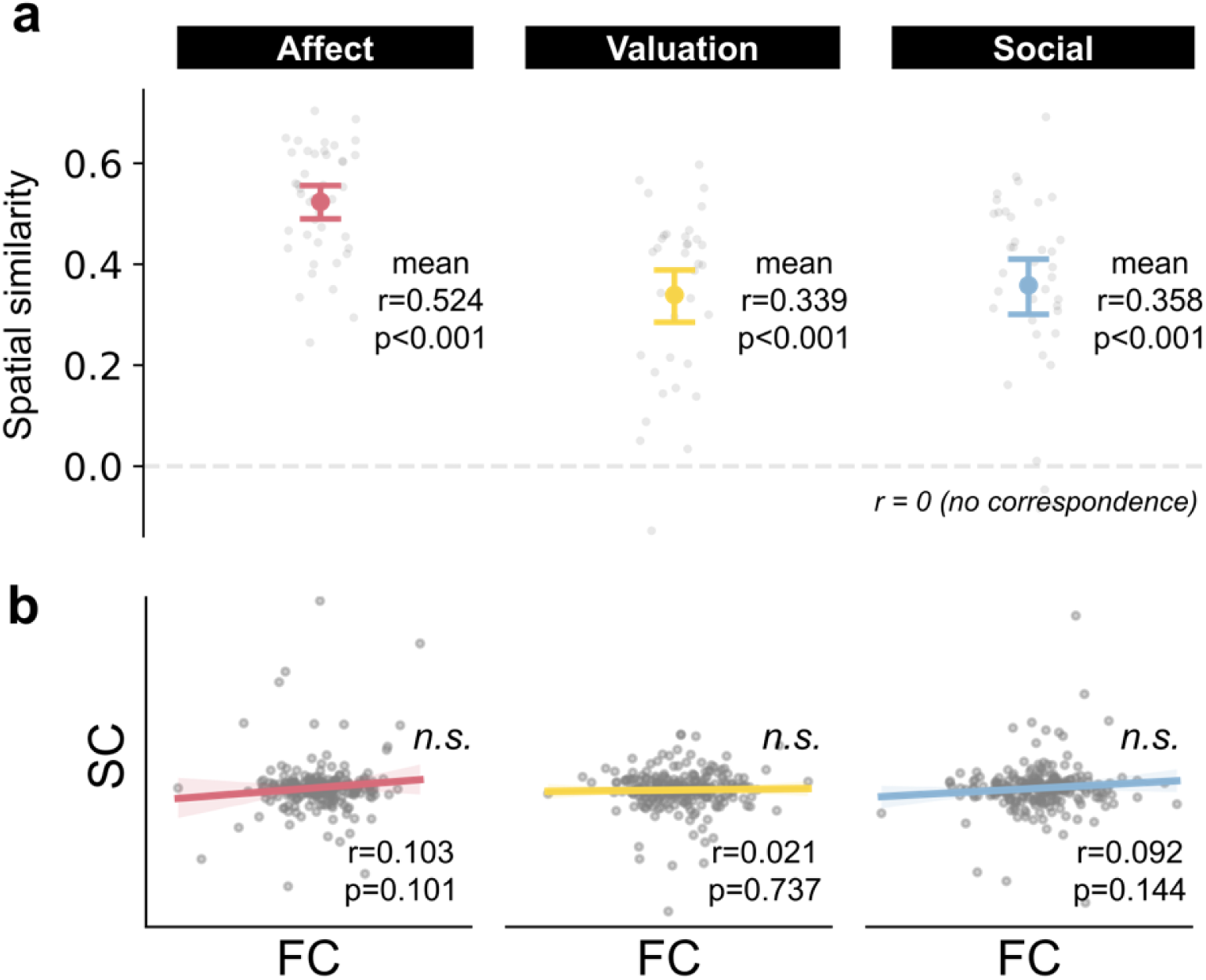
Quantitative comparison of structural-connectivity (SC) and functional-connectivity (FC) predictive models in Study 3. **a,** Spatial correspondence between SC- and FC-predicted activation maps within the VMPFC. For each participant, the voxel-wise Pearson correlation was computed between the activation map predicted by the SC model and that predicted by the FC model, separately for the Affect, Valuation, and Social domains. Each gray dot represents one participant. Colored symbols indicate the group mean ± 95% confidence interval. The dashed line denotes zero correspondence. Across all three domains, the majority of participants showed positive correlations, indicating that SC- and FC-based models recover broadly similar predicted functional topographies within the VMPFC (all mean *rs* > 0.33, all *ps* < 0.001). **b,** Correspondence between SC- and FC-derived regional regression weights. Each dot represents one brain region used in the connectivity-to-activity prediction model. Regression β-weights from the SC model are plotted against those from the FC model separately for each functional domain. Colored lines indicate least-squares regression fits. Correlations were weak across all three domains (all *rs* < 0.11, all *ps* > 0.10), indicating that SC and FC models rely on partially distinct connectivity features despite converging on similar predicted functional topographies.

## Reference

1. Delgado, M. R. et al. Viewpoints: Dialogues on the functional role of the ventromedial prefrontal cortex. Nat. Neurosci. 19, 1545–1552 (2016).

2. Damasio, H., Grabowski, T., Frank, R., Galaburda, A. M. & Damasio, A. R. The return of Phineas Gage: clues about the brain from the skull of a famous patient. Science 264, 1102–1105 (1994).

3. Gusnard, D. A. & Raichle, M. E. Searching for a baseline: Functional imaging and the resting human brain. Nat. Rev. Neurosci. 2, 685–694 (2001).

4. Hill, J. et al. Similar patterns of cortical expansion during human development and evolution. Proc. Natl. Acad. Sci. USA 107, 13135–13140 (2010).

5. Koban, L., Gianaros, P. J., Kober, H. & Wager, T. D. The self in context: brain systems linking mental and physical health. Nat. Rev. Neurosci. 22, 309–322 (2021).

6. Hiser, J. & Koenigs, M. The multifaceted role of the ventromedial prefrontal cortex in emotion, decision making, social cognition, and psychopathology. Biol. Psychiatry 83, 638–647 (2018).

7. Mackey, S. & Petrides, M. Architecture and morphology of the human ventromedial prefrontal cortex. Eur. J. Neurosci. 40, 2777–2796 (2014).

8. Mackey, S. & Petrides, M. Quantitative demonstration of comparable architectonic areas within the ventromedial and lateral orbital frontal cortex in the human and the macaque monkey brains. Eur. J. Neurosci. 32, 1940–1950 (2010).

9. Öngür, D., Ferry, A. T. & Price, J. L. Architectonic subdivision of the human orbital and medial prefrontal cortex. J. Comp. Neurol. 460, 425–449 (2003).

10. Lopez-Persem, A., Verhagen, L., Amiez, C., Petrides, M. & Sallet, J. The human ventromedial prefrontal cortex: sulcal morphology and its influence on functional organization. J. Neurosci. 39, 3627–3639 (2019).

11. Chang, L. J. et al. Endogenous variation in ventromedial prefrontal cortex state dynamics during naturalistic viewing reflects affective experience. Sci. Adv. 7, eabf7129 (2021).

12. Ojemann, J. G. et al. Anatomic localization and quantitative analysis of gradient refocused echo-planar fMRI susceptibility artifacts. NeuroImage 6, 156–167 (1997).

13. Weiskopf, N., Hutton, C., Josephs, O. & Deichmann, R. Optimal EPI parameters for reduction of susceptibility-induced BOLD sensitivity losses: A whole-brain analysis at 3 T and 1.5 T. NeuroImage 33, 493–504 (2006).

14. Murphy, K., Bodurka, J. & Bandettini, P. A. How long to scan? The relationship between fMRI temporal signal to noise ratio and necessary scan duration. NeuroImage 34, 565–574 (2007).

15. Jensen, K. T., Hennequin, G. & Mattar, M. G. A recurrent network model of planning explains hippocampal replay and human behavior. Nat. Neurosci. 27, 1340–1348 (2024).

16. Klein-Flügge, M. C., Bongioanni, A. & Rushworth, M. F. S. Medial and orbital frontal cortex in decision-making and flexible behavior. Neuron 110, 2743–2770 (2022).

17. Behrens, T. E. J. et al. Non-invasive mapping of connections between human thalamus and cortex using diffusion imaging. Nat. Neurosci. 6, 750–757 (2003).

18. Beckmann, M., Johansen-Berg, H. & Rushworth, M. F. S. Connectivity-based parcellation of human cingulate cortex and its relation to functional specialization. J. Neurosci. 29, 1175–1190 (2009).

19. Deen, B., Pitskel, N. B. & Pelphrey, K. A. Three systems of insular functional connectivity identified with cluster analysis. Cereb. Cortex 21, 1498–1506 (2011).

20. Tyszka, J. M. & Pauli, W. M. In vivo delineation of subdivisions of the human amygdaloid complex in a high-resolution group template. Hum. Brain Mapp. 37, 3979–3998 (2016).

21. Rudebeck, P. H., Bannerman, D. M. & Rushworth, M. F. S. The contribution of distinct subregions of the ventromedial frontal cortex to emotion, social behavior, and decision making. Cogn. Affect. Behav. Neurosci. 8, 485–497 (2008).

22. Roy, M., Shohamy, D. & Wager, T. D. Ventromedial prefrontal-subcortical systems and the generation of affective meaning. Trends Cogn. Sci. 16, 147–156 (2012).

23. Schurz, M., Radua, J., Aichhorn, M., Richlan, F. & Perner, J. Fractionating theory of mind: A meta-analysis of functional brain imaging studies. Neurosci. Biobehav. Rev. 42, 9–34 (2014).

24. Bartra, O., McGuire, J. T. & Kable, J. W. The valuation system: A coordinate-based meta-analysis of BOLD fMRI experiments examining neural correlates of subjective value. NeuroImage 76, 412–427 (2013).

25. Clithero, J. A. & Rangel, A. Informatic parcellation of the network involved in the computation of subjective value. Soc. Cogn. Affect. Neurosci. 9, 1289–1302 (2014).

26. Kober, H. et al. Functional grouping and cortical–subcortical interactions in emotion: A meta-analysis of neuroimaging studies. NeuroImage 42, 998–1031 (2008).

27. Laird, A. R. et al. Networks of task co-activations. NeuroImage 80, 505–514 (2013).

28. Laird, A. R., Lancaster, J. J. & Fox, P. T. BrainMap. Neuroinformatics 3, 65–77 (2005).

29. Fox, P. T. et al. Brainmap taxonomy of experimental design: Description and evaluation. Hum. Brain Mapp. 25, 185–198 (2005).

30. Barch, D. M. et al. Function in the human connectome: Task-fMRI and individual differences in behavior. NeuroImage 80, 169–189 (2013).

31. Ratan Murty, N. A., Bashivan, P., Abate, A., DiCarlo, J. J. & Kanwisher, N. Computational models of category-selective brain regions enable high-throughput tests of selectivity. Nat. Commun. 12, 5540 (2021).

32. Gu, Z. et al. NeuroGen: Activation optimized image synthesis for discovery neuroscience. NeuroImage 247, 118812 (2022).

33. Allen, E. J. et al. A massive 7T fMRI dataset to bridge cognitive neuroscience and artificial intelligence. Nat. Neurosci. 25, 116–126 (2022).

34. Saygin, Z. M. et al. Connectivity precedes function in the development of the visual word form area. Nat. Neurosci. 19, 1250–1255 (2016).

35. Osher, D. E. et al. Structural connectivity fingerprints predict cortical selectivity for multiple visual categories across cortex. Cereb. Cortex 26, 1668–1683 (2016).

36. Saygin, Z. M. et al. Anatomical connectivity patterns predict face selectivity in the fusiform gyrus. Nat. Neurosci. 15, 321–327 (2012).

37. Hiersche, K. J., Saygin, Z. M. & Osher, D. E. Connectivity and function are coupled across cognitive domains throughout the brain. Netw. Neurosci. 1–13 (2025).

38. Finn, E. S. et al. Functional connectome fingerprinting: identifying individuals using patterns of brain connectivity. Nat. Neurosci. 18, 1664–1671 (2015).

39. Gao, W. et al. Functional Network Development During the First Year: Relative Sequence and Socioeconomic Correlations. Cereb. Cortex 25, 2919–2928 (2015).

40. Fair, D. A. et al. Functional Brain Networks Develop from a “Local to Distributed” Organization. PLoS Comput. Biol. 5, e1000381 (2009).

41. Bzdok, D. et al. Segregation of the human medial prefrontal cortex in social cognition. Front. Hum. Neurosci. 7, (2013).

42. Eickhoff, S. B., Thirion, B., Varoquaux, G. & Bzdok, D. Connectivity-based parcellation: critique and implications. Hum. Brain Mapp. 36, 4771–4792 (2015).

43. Eickhoff, S. B., Yeo, B. T. T. & Genon, S. Imaging-based parcellations of the human brain. Nat. Rev. Neurosci. 19, 672–686 (2018).

44. Margulies, D. S. et al. Situating the default-mode network along a principal gradient of macroscale cortical organization. Proc. Natl. Acad. Sci. USA 113, 12574–12579 (2016).

45. Jackson, R. L., Bajada, C. J., Lambon Ralph, M. A. & Cloutman, L. L. The graded change in connectivity across the ventromedial prefrontal cortex reveals distinct subregions. Cereb. Cortex 30, 165–180 (2020).

46. Casamitjana, A. et al. A probabilistic histological atlas of the human brain for MRI segmentation. Nature 648, 678–685 (2025).

47. Lockwood, P. L., Apps, M. A. J. & Chang, S. W. C. Is there a ‘social’ brain? Implementations and algorithms. Trends Cogn. Sci. 24, 802–813 (2020).

48. Koster-Hale, J. & Saxe, R. Theory of mind: a neural prediction problem. Neuron 79, 836–848 (2013).

49. Mahmoodi, A. & Rushworth, M. F. S. Computational origins of cortical brain circuits for social cognition. Nat. Rev. Neurosci. 27, 345–356 (2026).

50. Eichenbaum, H. Prefrontal–hippocampal interactions in episodic memory. Nat. Rev. Neurosci. 18, 547–558 (2017).

51. Bechara, A., Damasio, H., Tranel, D. & Damasio, A. R. Deciding advantageously before knowing the advantageous strategy. Science 275, 1293–1295 (1997).

52. Carmichael, S. T. & Price, J. L. Architectonic subdivision of the orbital and medial prefrontal cortex in the macaque monkey. J. Comp. Neurol. 346, 366–402 (1994).

53. Roberts, A. C. & Clarke, H. F. Why we need nonhuman primates to study the role of ventromedial prefrontal cortex in the regulation of threat- and reward-elicited responses. Proc. Natl. Acad. Sci. 116, 26297–26304 (2019).

54. Barbas, H. General cortical and special prefrontal connections: principles from structure to function. Annu. Rev. Neurosci. 38, 269–289 (2015).

55. Valk, S. L. et al. Shaping brain structure: Genetic and phylogenetic axes of macroscale organization of cortical thickness. Sci. Adv. 6, (2020).

56. Guerrero, J. M., Cabrera, A. J. & Vasquez, J. C. Evolution and functions of the cerebral cortex – a review. Preprint at 10.20944/preprints202310.1210.v1 (2023).

57. Goulas, A., Margulies, D. S., Bezgin, G. & Hilgetag, C. C. The architecture of mammalian cortical connectomes in light of the theory of the dual origin of the cerebral cortex. Cortex 118, 244–261 (2019).

58. Benoit, R. G. & Schacter, D. L. Specifying the core network supporting episodic simulation and episodic memory by activation likelihood estimation. Neuropsychologia 75, 450–457 (2015).

59. Lieberman, M. D., Straccia, M. A., Meyer, M. L., Du, M. & Tan, K. M. Social, self, (situational), and affective processes in medial prefrontal cortex (MPFC): Causal, multivariate, and reverse inference evidence. Neurosci. Biobehav. Rev. 99, 311–328 (2019).

60. Rolls, E. T. The hippocampus, ventromedial prefrontal cortex, and episodic and semantic memory. Prog. Neurobiol. 217, 102334 (2022).

61. Doeller, C. F., Barry, C. & Burgess, N. Evidence for grid cells in a human memory network. Nature 463, 657–661 (2010).

62. Critchley, H. D. et al. Human cingulate cortex and autonomic control: converging neuroimaging and clinical evidence. Brain 126, 2139–2152 (2003).

63. Naqvi, N., Tranel, D. & Bechara, A. Visceral and decision-making functions of the ventromedial prefrontal cortex. in The Orbitofrontal Cortex (eds Zald, D. & Rauch, S.) 325–354 (Oxford University Press, 2006).

64. Foster, B. L. et al. A tripartite view of the posterior cingulate cortex. Nat. Rev. Neurosci. 24, 173–189 (2023).

65. Gratton, C. & Braga, R. M. Dense phenotyping of human brain network organization using precision fMRI. Annu. Rev. Psychol. (in the press).

66. Chiavaras, M. M. & Petrides, M. Orbitofrontal sulci of the human and macaque monkey brain. J. Comp. Neurol. 422, 35–54 (2000).

67. Ahn, W.-Y., Haines, N. & Zhang, L. Revealing neurocomputational mechanisms of reinforcement learning and decision-making with the hBayesDM package. *Comput*. Psychiatry 1, 24–57 (2017).

68. Karvelis, P., Paulus, M. P. & Diaconescu, A. O. Individual differences in computational psychiatry: A review of current challenges. Neurosci. Biobehav. Rev. 148, 105137 (2023).

69. Kahnt, T., Chang, L. J., Park, S. Q., Heinzle, J. & Haynes, J.-D. Connectivity-Based Parcellation of the Human Orbitofrontal Cortex. J. Neurosci. 32, 6240–6250 (2012).

70. Chase, H. W., Grace, A. A., Fox, P. T., Phillips, M. L. & Eickhoff, S. B. Functional differentiation in the human ventromedial frontal lobe: A data-driven parcellation. Hum. Brain Mapp. 41, 3266–3283 (2020).

71. Yarkoni, T., Poldrack, R. A., Nichols, T. E., Van Essen, D. C. & Wager, T. D. Large-scale automated synthesis of human functional neuroimaging data. Nat. Methods 8, 665–670 (2011).

72. Zhang, Y., et al. Qwen3 Embedding: Advancing Text Embedding and Reranking Through Foundation Models. Preprint at 10.48550/arXiv.2506.05176 (2025).

73. Chang, L. J., Yarkoni, T., Khaw, M. W. & Sanfey, A. G. Decoding the Role of the Insula in Human Cognition: Functional Parcellation and Large-Scale Reverse Inference. Cereb. Cortex 23, 739–749 (2013).

74. Tovar, D. T. & Chavez, R. S. Large-scale functional coactivation patterns reflect the structural connectivity of the medial prefrontal cortex. Soc. Cogn. Affect. Neurosci. 16, 875–882 (2021).

75. Eickhoff, S. B. et al. Co-activation patterns distinguish cortical modules, their connectivity and functional differentiation. NeuroImage 57, 938–949 (2011).

76. Reuter, N. et al. CBPtools: a Python package for regional connectivity-based parcellation. Brain Struct. Funct. 225, 1261–1275 (2020).

77. Van Essen, D. C. et al. The WU-Minn Human Connectome Project: An overview. NeuroImage 80, 62–79 (2013).

78. Glasser, M. F. et al. A multi-modal parcellation of human cerebral cortex. Nature 536, 171–178 (2016).

79. Wang, Y. et al. Multimodal mapping of the face connectome. *Nat*. Hum. Behav. 4, 397–411 (2020).

80. Khosla, M., Ngo, G. H., Jamison, K., Kuceyeski, A. & Sabuncu, M. R. Cortical response to naturalistic stimuli is largely predictable with deep neural networks. Sci. Adv. 7, eabe7547 (2021).

81. LeBel, A. et al. A natural language fMRI dataset for voxelwise encoding models. Sci. Data 10, 555 (2023).

82. Deichmann, R., Gottfried, J. A., Hutton, C. & Turner, R. Optimized EPI for fMRI studies of the orbitofrontal cortex. NeuroImage 19, 430–441 (2003).

83. Volz, S., Callaghan, M. F., Josephs, O. & Weiskopf, N. Maximising BOLD sensitivity through automated EPI protocol optimisation. NeuroImage 189, 159–170 (2019).

84. Olman, C. A., Davachi, L. & Inati, S. Distortion and Signal Loss in Medial Temporal Lobe. PLoS ONE 4, e8160 (2009).

85. Constantinescu, A. O., O’Reilly, J. X. & Behrens, T. E. J. Organizing conceptual knowledge in humans with a gridlike code. Science 352, 1464–1468 (2016).

86. Baram, A. B., Muller, T. H., Nili, H., Garvert, M. M. & Behrens, T. E. J. Entorhinal and ventromedial prefrontal cortices abstract and generalize the structure of reinforcement learning problems. Neuron 109, 713–723.e7 (2021).

87. Fan, L. et al. The human brainnetome atlas: a new brain atlas based on connectional architecture. Cereb. Cortex 26, 3508–3526 (2016).

88. King, M., Hernandez-Castillo, C. R., Poldrack, R. A., Ivry, R. B. & Diedrichsen, J. Functional boundaries in the human cerebellum revealed by a multi-domain task battery. Nat. Neurosci. 22, 1371–1378 (2019).

89. Harms, M. P. et al. Extending the Human Connectome Project across ages: Imaging protocols for the Lifespan Development and Aging projects. NeuroImage 183, 972–984 (2018).

90. Glasser, M. F. et al. The minimal preprocessing pipelines for the Human Connectome Project. NeuroImage 80, 105–124 (2013).

91. Somerville, L. H. et al. The lifespan human connectome project in development: a large-scale study of brain connectivity development in 5–21 year olds. NeuroImage 183, 456–468 (2018).

92. Bookheimer, S. Y. et al. The Lifespan Human Connectome Project in Aging: An overview. NeuroImage 185, 335–348 (2019).

93. Milham, M. P. et al. An open resource for non-human primate imaging. Neuron 100, 61–74.e2 (2018).

94. Xu, T. et al. Cross-species functional alignment reveals evolutionary hierarchy within the connectome. NeuroImage 223, 117346 (2020).

95. Samara, Z. et al. Human orbital and anterior medial prefrontal cortex: intrinsic connectivity parcellation and functional organization. Brain Struct. Funct. 222, 2941–2960 (2017).

96. Craddock, R. C., James, G. A., Holtzheimer, P. E., Hu, X. P. & Mayberg, H. S. A whole brain fMRI atlas generated via spatially constrained spectral clustering. Hum. Brain Mapp. 33, 1914–1928 (2012).

97. Shen, X., Tokoglu, F., Papademetris, X. & Constable, R. T. Groupwise whole-brain parcellation from resting-state fMRI data for network node identification. NeuroImage 82, 403–415 (2013).

98. Schaefer, A. et al. Local-global parcellation of the human cerebral cortex from intrinsic functional connectivity MRI. Cereb. Cortex 28, 3095–3114 (2018).

99. Neubert, F.-X., Mars, R. B., Sallet, J. & Rushworth, M. F. S. Connectivity reveals relationship of brain areas for reward-guided learning and decision making in human and monkey frontal cortex. Proc. Natl. Acad. Sci. USA 112, E2695–E2704 (2015).

100. de La Vega, A., Chang, L. J., Banich, M. T., Wager, T. D. & Yarkoni, T. Large-scale meta-analysis of human medial frontal cortex reveals tripartite functional organization. J. Neurosci. 36, 6553–6562 (2016).

101. Liu, H., Qin, W., Qi, H., Jiang, T. & Yu, C. Parcellation of the human orbitofrontal cortex based on gray matter volume covariance. Hum. Brain Mapp. 36, 538–548 (2015).

102. Morris, L. S. et al. Ultra-high field MRI reveals mood-related circuit disturbances in depression: a comparison between 3-Tesla and 7-Tesla. Transl. Psychiatry 9, 94 (2019).

## Supplementary References

Ashburner, J., & Friston, K. J. (2000). Voxel-based morphometry—The Methods. NeuroImage, 11(6), 805–821. 10.1006/nimg.2000.0582

Delgado, M. R., Beer, J. S., Fellows, L. K., Huettel, S. A., Platt, M. L., Quirk, G. J., & Schiller, D. (2016). Viewpoints: Dialogues on the functional role of the ventromedial prefrontal cortex. Nature Neuroscience, 19(12), 1545–1552. 10.1038/nn.4438

Eickhoff, S. B., Thirion, B., Varoquaux, G., & Bzdok, D. (2015). Connectivity-based parcellation: Critique and implications. Human Brain Mapping, 36(12), 4771–4792. 10.1002/hbm.22933

Eickhoff, S. B., Yeo, B. T. T., & Genon, S. (2018). Imaging-based parcellations of the human brain. Nature Reviews Neuroscience, 19(11), 672–686. 10.1038/s41583-018-0071-7

Hansen, J. Y., Shafiei, G., Markello, R. D., Smart, K., Cox, S. M. L., Nørgaard, M., Beliveau, V., Wu, Y., Gallezot, J.-D., Aumont, É., Servaes, S., Scala, S. G., DuBois, J. M., Wainstein, G., Bezgin, G., Funck, T., Schmitz, T. W., Spreng, R. N., Galovic, M., … Misic, B. (2022). Mapping neurotransmitter systems to the structural and functional organization of the human neocortex. Nature Neuroscience, 25(11), 1569–1581. 10.1038/s41593-022-01186-3

Hiser, J., & Koenigs, M. (2018). The multifaceted role of the ventromedial prefrontal cortex in emotion, decision making, social cognition, and psychopathology. Biological Psychiatry, 83(8), 638–647. 10.1016/j.biopsych.2017.10.030

Milham, M. P., Ai, L., Koo, B., Xu, T., Amiez, C., Balezeau, F., Baxter, M. G., Blezer, E. L. A., Brochier, T., Chen, A., Croxson, P. L., Damatac, C. G., Dehaene, S., Everling, S., Fair, D. A., Fleysher, L., Freiwald, W., Froudist-Walsh, S., Griffiths, T. D., … Schroeder, C. E. (2018). An open resource for non-human primate imaging. Neuron, 100(1), 61–74.e2. 10.1016/j.neuron.2018.08.039

Markello, R. D., Hansen, J. Y., Liu, Z. Q., Bazinet, V., Shafiei, G., Suárez, L. E., Blostein, N., Seidlitz, J., Baillet, S., Satterthwaite, T. D., Chakravarty, M. M., Raznahan, A., & Misic, B. (2022). Neuromaps: structural and functional interpretation of brain maps. Nature Methods, 19(11), 1472–1479.

Rudebeck, P. H., Bannerman, D. M., & Rushworth, M. F. S. (2008). The contribution of distinct subregions of the ventromedial frontal cortex to emotion, social behavior, and decision making. *Cognitive, Affective*, & Behavioral Neuroscience, 8(4), 485–497. 10.3758/CABN.8.4.485

Schaefer, A., Kong, R., Gordon, E. M., Laumann, T. O., Zuo, X.-N., Holmes, A. J., Eickhoff, S. B., & Yeo, B. T. T. (2018). Local-global parcellation of the human cerebral cortex from intrinsic functional connectivity MRI. Cerebral Cortex, 28(9), 3095–3114. 10.1093/cercor/bhx179

Wang, Z., Li, Y., Childress, A. R., & Detre, J. A. (2014). Brain entropy mapping using fMRI. PLOS ONE, 9(3), e89948. 10.1371/journal.pone.0089948

